# Time series transcriptomics reveals a *BBX32*-directed control of dynamic acclimation to high light in mature *Arabidopsis* leaves

**DOI:** 10.1101/2020.12.23.424212

**Authors:** Ruben Alvarez-Fernandez, Christopher A. Penfold, Gregorio Galvez-Valdivieso, Marino Exposito-Rodriguez, Ellie J. Stallard, Laura Bowden, Jonathan D. Moore, Andrew Mead, Phillip A. Davey, Jack S.A. Matthews, Jim Beynon, Vicky Buchanan-Wollaston, David L. Wild, Tracy Lawson, Ulrike Bechtold, Katherine Denby, Philip M. Mullineaux

## Abstract

The photosynthetic capacity of mature leaves increases after several days’ exposure to constant or intermittent episodes of high light (HL) and is manifested primarily as changes in chloroplast physiology. This is termed dynamic acclimation but how it is initiated and controlled is unknown. From fully expanded Arabidopsis leaves, we determined HL-dependent changes in transcript abundance of 3844 genes in a 0-6h time-series transcriptomics experiment. It was hypothesised that among such genes were those that contribute to the initiation of dynamic acclimation. By focussing on HL differentially expressed transcription (co-)factor (TF) genes and applying dynamic statistical modelling to the temporal transcriptomics data, a gene regulatory network (GRN) of 47 predominantly photoreceptor-regulated (co)-TF genes was inferred. The most connected gene in this network was *B-BOX DOMAIN CONTAINING PROTEIN32* (*BBX32*). Plants over-expressing *BBX32* were strongly impaired in dynamic acclimation and displayed perturbed expression of genes involved in its initiation. These observations led to demonstrating that as well as regulation of dynamic acclimation by *BBX32*, *CRYPTOCHROME1, LONG HYPOCOTYL5*, *CONSTITUTIVELY PHOTOMORPHOGENIC1* and *SUPPRESSOR OF PHYA-105* are also important regulators of this process. Additionally, the *BBX32*-centric GRN provides a view of the transcriptional control of dynamic acclimation distinct from other photoreceptor-regulated processes, such as seedling photomorphogenesis.

---

The exposure of plants to increased light intensities can lead to the development of enhanced photosynthetic capacity in mature leaves and is termed dynamic acclimation. Dynamic acclimation, an important determinant of plant fitness or crop yield, is under genetic as well as environmental control and includes changes in the expression of many genes (Murchie and Horton, 1997; Walters et al., 1999; Oguchi et al., 2003; Murchie et al., 2005; Eberhard et al., 2008; Athanasiou et al., 2010; Schottler and Toth, 2014; van Rooijen et al 2015; Vialet-Chabrand et al., 2017).

In young expanding leaves, acclimation to increased light intensity (hereafter termed high light; HL) brings about increased photosynthetic capacity by eliciting changes in both leaf morphology, such as altered leaf and vascular diameter in minor veins (Oguchi et al., 2003; Terashima et al., 2011; Adams III et al., 2014; Vialet-Chabrand et al., 2017) and changes to chloroplast physiology such as adjustments to the composition of reaction centers and light harvesting antennae (Walters et al., 1999, Murchie and Horton, 1997; Murchie et al., 2005; Drozak and Romanowska, 2006; Vialet-Chabrand et al., 2017). In contrast, in mature leaves exposure to sustained or episodic HL brings about changes primarily in chloroplast physiology that raise the light use efficiency of photosynthesis, which can reflect increased rates of photosynthesis and/or a decreased number of photosystem II (PSII) reaction centers (Walters et al., 1999; Murchie et al., 2005; Athanasiou et al., 2010; van Rooijen et al., 2015; Vialet-Chabrand et al., 2017).

It is not known how HL exposure initiates chloroplast-level dynamic acclimation. However, an important lead is provided from an early study (Walters et al., 1999). This was a comparison of Arabidopsis photoreceptor signalling mutants’ photosynthetic capacity and PSII efficiency grown under two different light intensities (PPFDs) of 100 and 400 µmol m^-2^ s^-1^ (PPFD stands for photosynthetically active photon flux density). From this study, it was proposed that a PHYTOCHROMEA (PHYA), PHYB and CRYPTOCHROME1 (CRY1) photoreceptor driven CONSTITUTIVELY PHOTOMORPHOGENIC/DE-ETIOLATED1/FUSCA (COP/DET/FUS) regulatory module could transmit signals from the nucleus to chloroplasts to participate in establishing increased photosynthetic capacity (Walters et al.,1999). In support of this, photosynthesis-associated nuclear genes (PhANGs) are among the most enriched gene classes subject to control from photoreceptors in de-etiolating seedlings (Chory and Peto, 1990; Holtan et al., 2011; Shikata et al., 2014; Li et al., 2015; Ganguly et al., 2015; Pham et al., 2018). Various combinations of the 11 *COP/DET/FUS* loci (Lau and Deng, 2012), in conjunction with other regulatory genes, control the integration of signals from photoreceptors and are central to many plant-light environment interactions including seedling photomorphogenesis, the shade avoidance response, stomatal opening and development, the timing of flowering and cross-talk between phytohormone and light signalling (Dong et al., 2014; Huang et al., 2014; Lau and Deng, 2012; Pham et al., 2018).

The establishment of dynamic acclimation takes up to 6 days (Athanasiou et al., 2010) and prior to this, plants must deal with HL by dissipating excitation energy to minimise irreversible photoinhibition. Photoinhibition is caused by oxidative damage to the photosynthetic apparatus brought about by increased production of singlet oxygen (^1^O_2_; Triantaphyllides et al., 2008; Ramel et al., 2013; Mullineaux et al., 2018). This is achieved by a combination of non-photochemical and photochemical quenching (NPQ and PQ respectively; Baker, 2008; Eberhard et al., 2008; Ruban and Belgio, 2014). All leaves have an extant capability to dissipate excitation energy, which is augmented under HL by localised and systemic induction of further photo-protective NPQ and/or PQ (Long et al., 1994; Karpinski et al., 1999; Eberhard et al., 2008; Li et al., 2009; Galvez-Valdivieso et al., 2009; Suorsa et al., 2012; Ruban and Belgio, 2014). NPQ is the resonance transfer of excitation energy to xanthophyll carotenoids from excited chlorophylls and its subsequent thermal dissipation (Li et al., 2000; Baker, 2008). PQ is the dissipation of excitation energy by consumption of reducing equivalents by a range of metabolic processes including enhanced photosynthetic capacity, but is also associated with increased foliar levels of hydrogen peroxide (H_2_O_2_) and the superoxide anion (Long et al., 1994; Wingler et al., 2000; Badger et al., 2000; Schiebe, 2004; Schieble et al., 2004; Streb et al., 2005; Diaz et al., 2007; Eberhard et al., 2008; Driever and Baker, 2010; Kangasjarvi et al., 2012; Schiebe and Dietz, 2012; Lawson et al., 2013; Heyno et al., 2014; Mullineaux et al., 2018).

In this study, we hypothesised that the first hours of exposure of mature leaves to HL initiates dynamic acclimation although an increase in photosynthetic capacity takes several days to develop. This hypothesis is an extension of an earlier proposal regarding the temporal order of events leading to protection against oxidative stress-induced photoinhibition, the re-structuring of light harvesting antennae and PSI and PSII reaction centers (Eberhard et al., 2008). We tested this hypothesis by first developing HL and low light (LL) time series transcriptomics datasets from a defined fully expanded leaf of LL-grown Arabidopsis plants. Analysis of these data revealed that processes interpreted as the first steps in establishing dynamic acclimation could be identified and were largely complete within 4h of HL exposure. From comparison with other published transcriptomics datasets, we noted an over-representation of differentially expressed transcription (co)factor genes involved in light- and photoreceptor-regulated photomorphogenesis of seedlings. This led us via dynamic statistical modelling to identify *B-BOX DOMAIN CONTAINING PROTEIN32* (*BBX32*) as a negative regulator of dynamic acclimation. *BBX32* encodes a Zn-finger transcription co-factor which forms associations with diverse transcription regulators, such as LONG HYPOCOTYL5 (HY5) and BBX4 to alter their function and is most studied in the context of seedling photomorphogenesis and the control of flowering time (Holtan et al., 2011; Park et al., 2011; Gangappa and Botto, 2014; Tripathy et al., 2017). In turn, the involvement of *BBX32* led us to identify *HY5*, *CRY1*, *PHYTOCHROME INTERACTING FACTOR* (*PIF*) genes and *COP1/SUPPRESSOR OF phytochromeA-105* (*SPA*) genes as further regulators of dynamic acclimation. It was concluded that many signalling aspects of photoreceptor-driven seedling photomorphogenesis are also active in mature fully expanded leaves to control dynamic acclimation.

## Results

### GO analysis of time series transcriptomics of HL-exposed leaves provides insights into the initiation of dynamic acclimation

The starting point for this study was the development of a HL time series transcriptomics experiment. Our plan was to subject groups of time-resolved differentially expressed genes (DEGs) to Variational Bayesian State Space Modelling (VBSSM; see Methods), which requires highly resolved time series data (Beal et al., 2005; Penfold and Wild, 2011; Penfold and Buchanan-Wollaston, 2014; Bechtold et al., 2016). Therefore, we opted for 30 min sampling over a 6h HL period beginning 1h after subjective dawn (see Methods). We chose this time period because it spans the initiation of both the short term and long-term acclimation to HL proposed by Eberhard et al. (2008; see Introduction).

Full transcriptome profiles using CATMA microarrays (Sclep et al., 2007) were obtained from leaf 7 of HL-exposed plants along with parallel LL controls (see Methods). Microarray analysis of variance (MAANOVA; Wu et al., 2003; see Methods) was used to extract expression values from each probe for every treatment for each technical and biological replicate. To determine DEGs that showed a significant difference between HL-exposed leaves and the LL controls over all or part of the time period, a Gaussian process two-sample test (GP2S; Stegle et al., 2010) was applied and 4069 probes were selected with a Bayes factor score >10 which corresponded to 3844 DEGs (Supplemental Data Set 1). The full data set is deposited with Gene Expression Omnibus (GEO; GSE78251).

To obtain further insight into the overall response to HL at the molecular level, hierarchical co-cluster analysis of the 3844 DEGs was carried out using SplineCluster (Heard et al., 2005). We reasoned that groups of DEGs that display similar temporal patterns of expression could be co-regulated and clustering would be useful in identifying groups of genes for dynamic modelling. On the basis of comparing temporal gene expression patterns in both the HL-exposed and control LL leaves (see Methods), the 3844 DEGs were divided into 43 temporal clusters (Fig. 1A; Supplemental Data Set 1). The outcome of this co-clustering was differential transcript abundance between LL and HL conditions superimposed on a range of temporal expression trends across 6h of the diel. Plotted examples showing a range of temporal and differential expression in 6 clusters can be viewed in Supplemental Figure 1.

**Figure 1.**
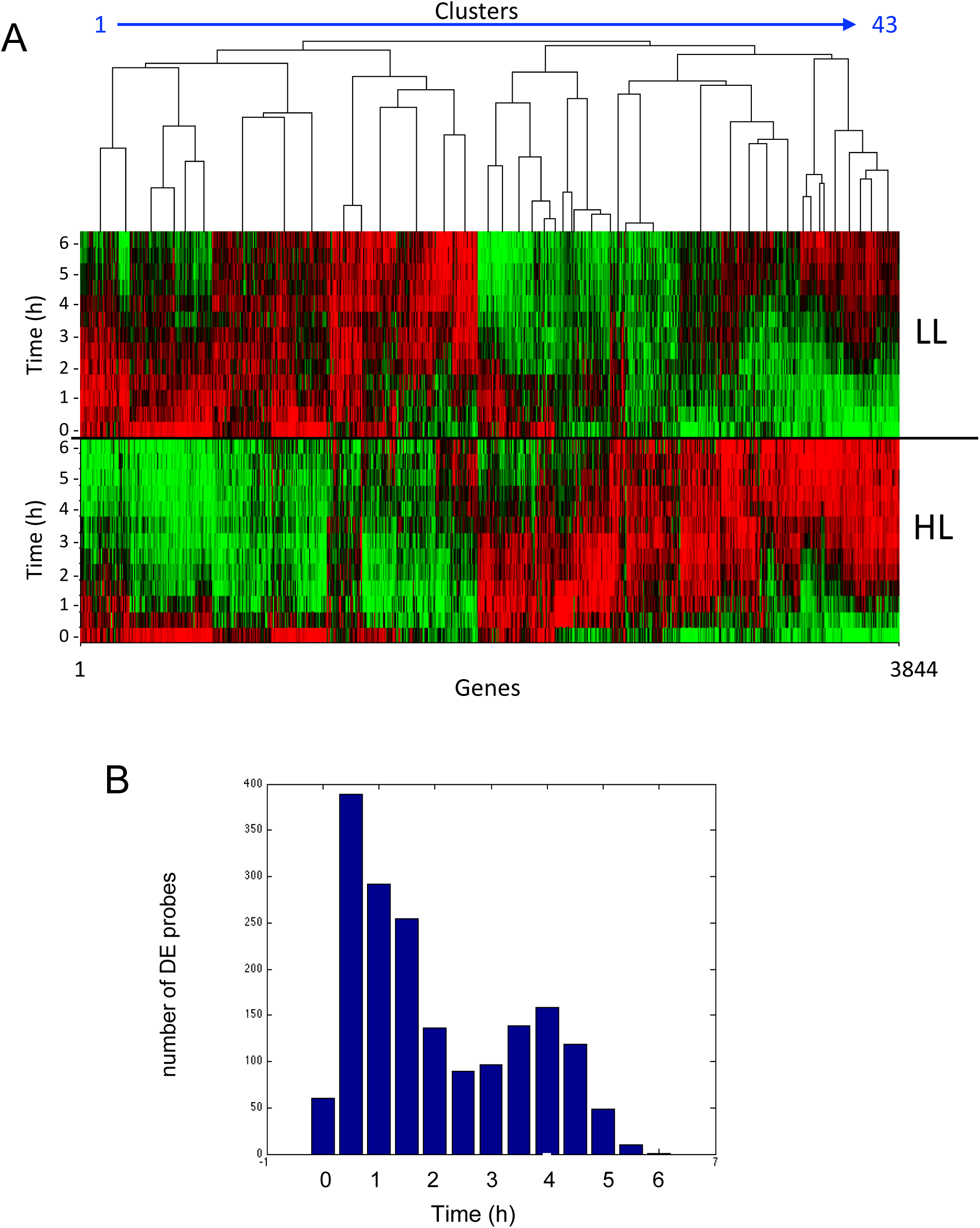
**Temporal patterns of gene expression in LL- and HL-exposed leaf 7.** **A.** Visual output of co-clustered expression values by SplineCluster. This was done for the 3844 genes already identified as differentially expressed in HL vs LL over the time of the experiment (see Results and Supplemental Dataset 1). The values range from log2 2.5 (red) to –log2 2.0 (green). The 43 temporal clusters can be counted in the accompanying dendrogram. The timepoints are shown on the y-axes for the HL and LL gene expression. **B.** The number of HL/LL differentially expressed probes first appearing at each time point.

The clusters are ordered such that 1-13 show general transcript abundance to be lower in HL vs LL samples and/or displayed a downward pattern over the diel (Fig.1A; e.g. clusters 3 and 13 in Supplemental Fig. 1). This pattern changes progressively with increasing degree of expression being higher in HL than LL but against a descending diel pattern in clusters 14-20 (Fig. 1A, e.g. cluster 18 in Supplemental Fig. 1), followed by transient but progressively increasingly greater differential transcript levels in HL samples compared with LL in clusters 21-26 (Fig. 1A, e.g. cluster 23 in Supplemental Fig. 1) to progressively sustained periods of higher expression in HL compared with LL from cluster 27-43 against a background of level or increasing transcript levels across the diel (Fig. 1A; e.g. clusters 33 and 43 in Supplemental Figure 1).

To gain a better view of the timings of differential expression in response to HL, the DEGs from the time-local GP2S (Supplemental Dataset 1) were used to identify intervals of differential expression as described by Windram et al. (2012). A histogram of the time of first differential expression (HL compared to LL) is shown in Figure 1B and indicates that response to HL is rapid with >700 genes becoming differentially expressed by 1 hour into the HL time course. Nevertheless, it was also clear that changes in transcript abundance were being initiated for significant numbers of genes up to 4h HL. In summary, the response of the leaf 7 transcriptome to HL entails changed expression in response to the stimulus, with changes occurring across the time of the experiment against a backdrop of complex changes in transcript abundance across 6h out of an 8h photoperiod.

Gene Ontology (GO) analysis showed that clusters 22, 23, 25 were highly enriched for generic abiotic stress-defensive genes (P value ≤ 0.1, Bonferroni corrected; Supplemental Data Set 2). In contrast, some of the other clusters displayed a different set of GO function enrichments (Supplemental Data Set 2). These multiple enriched sets were consistent with a re-adjustment to cellular metabolism. For example, in clusters 39 and 41 – 43 with generally higher expression in HL compared with LL, there was over-representation of genes associated with amino acid and protein synthesis respectively. Among the clusters showing a lowered expression in HL compared with LL, there was enrichment for genes associated with cell wall metabolism (callose deposition, cell wall thickening Cluster 1), phenylpropanoid and glucosinolate metabolism (Clusters 1 and 10 respectively), basal resistance to infection (Cluster 3) and chromatin re-modelling (cluster 10).

### Induction of dynamic acclimation

To test our interpretation of the HL time series data, we determined if dynamic acclimation could be induced by exposing a plant every day to 4h HL (see Methods). This period of HL exposure was chosen since most differential expression had been initiated by this time (Fig. 1B). Other than being shortened to 4h, the environmental conditions were the same as for the time series transcriptomics experiment (see Methods). The daily HL regime brought about a stepwise increase in the operating efficiency of PSII (Fq’/Fm’) (Baker, 2008) of fully expanded leaves (Fig. 2A; Supplemental Data Set 3). By day 5, PSII operating efficiency had increased substantially (e.g. 78% at 800 µmol m^-2^ s^-1^ actinic PPFD; Fig. 2B; see also Fig.4B). This pattern was followed by equivalent changes in Fv’/Fm’ and Fq’/Fv’ (Supplemental Fig. 2A; Supplemental Data Set 3). Fv’/Fm’ indicates the maximum operating efficiency of PSII at a given PPFD and a rise in this parameter indicates a decline in NPQ (Baker, 2008). Fq’/Fv’ is the PSII efficiency factor and is mathematically identical to the coefficient for PQ (*q_p_*) and indicates increased capacity to drive electron transport (Baker, 2008). Control LL-grown plants of the same age as the plants subjected to 5 daily HL treatments did not show these changes in chlorophyll fluorescence (CF) parameters (Supplemental Fig. 2B; Supplemental Data Set 3).

**Figure 2.**
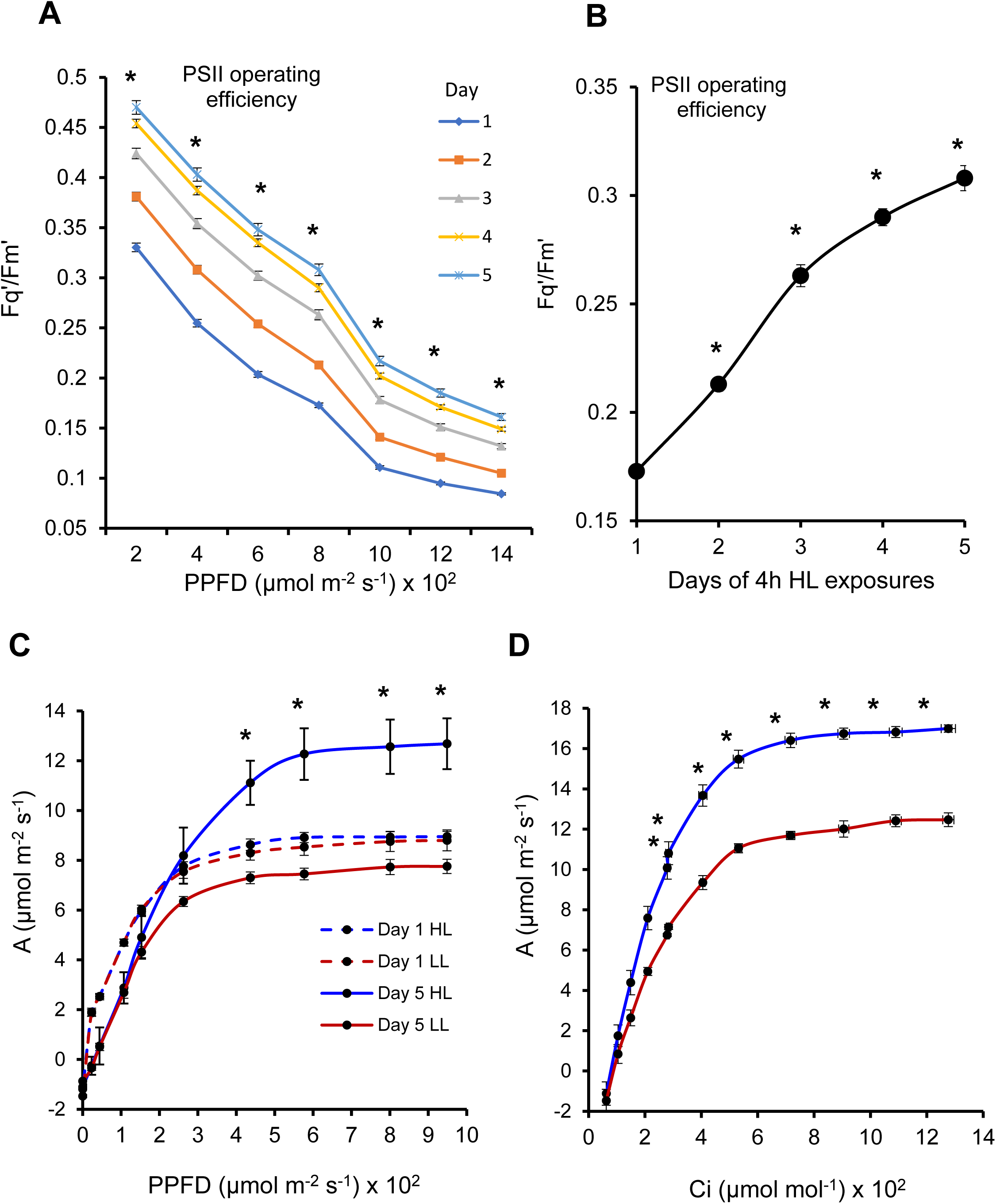
**Induction of dynamic acclimation by repeated daily exposure to HL.** **(A)** Plants were exposed daily to 4h HL and Fq’/Fm’ determined for mature leaves. After the HL, plants were dark adapted and imaged under increasing actinic PPFD from 200–to- 1400 µmol m^-2^ s^-1^ in 200 µmol m^-2^ s^-1^ increments every 5 min. The data were collected as CF images and processed digitally to collect values from mature leaves. The plants were treated in this way daily for 5 days: day 1 (blue), day 2 (red), day 3 (olive green), day 4 (purple) and day 5 (light blue). The data (mean ± SE) correspond to 38 plants at 24 – 28 dpg over 6 experiments and the asterisks show differences in CF parameters between days 1 and 5 were significant (P ≤ 0.001; ANOVA and TukeyHSD). The full statistical data comparing all days of HL exposure are provided in Supplemental Data Set 5. **(B)** Daily changes in Fq’/Fm’ plotted from the data in A and Supplemental Data Set 5 (right panel). Fq’/Fm’ values are from the same plants over the daily HL exposures showing the the increase in PSII operating efficiency at 800 µmol m^-2^ s^-1^ PPFD actinic light over the 5 days of the experiments. **(C)** Photosynthesis plotted as CO_2_ assimilation rate (*A*) as a function of actinic PPFD in mature leaf 7 (mean ± SE; n = 8 plants for each treatment; 49 dpg). Measurements were taken the day after 1 (dashed lines) and 5 days (solid lines) of daily 4h HL exposures (blue lines) along with the LL control plants (red lines) not subjected to this treatment. **(D)** Photosynthesis plotted as CO_2_ assimilation rate (*A*) as function of leaf internal CO_2_ concentration (Ci) in mature leaf 7 (mean ± SE; n = 8 plants for each treatment; 49 dpg). Measurements were taken the day after 5 days of daily 4h HL exposures (blue line) along with the LL control (red line). *A* was determined by Infra-Red Gas Analysis (see Methods). Asterisks indicate significant differences (P < 0.02; covariant T and two-tailed F tests) between LL and HL-exposed plants.

The first exposure to HL (day 1) did not result in irreversible photoinhibition (Supplemental Fig. 3A) or significant tissue damage (Supplemental Fig. 3B). This was confirmed in the HL time series data, which used the same PPFDs, in which steady levels of transcripts for genes considered to be markers for H_2_O_2_ (*APX2* and *FER1*, Ball et al., 2004; Gadjev et al., 2006) rose but those associated with ^1^O_2_ – induced signalling (*AAA-ATPase* and *BAP1*, Ramel et al., 2013) remained unchanged or declined (Supplemental Fig. 3C). The changes in the expression of these marker genes indicated the HL treatment used in the time series transcriptomics experiments, also did not elicit photodamage and provided conditions that could promote dynamic acclimation.

The increased operating efficiency of PSII (Fq’/Fm’ and also Fq’/Fv’) after the 5-day HL treatment (Fig. 2A; Supplemental Fig. 2A; Supplemental Data Set 3) could have reflected enhanced photosynthetic capacity. To test this possibility, gas-exchange measurements for photosynthesis were carried out (see Methods). The same experiment was repeated and, in the photoperiod following the last HL treatment, measurements of CO_2_ assimilation rates (*A*) over a range of light intensities in fully expanded leaf 7 of these plants (Boyes et al., 1998) were taken. This showed that the light-saturated photosynthetic rate (*A*_sat_) was significantly greater (P < 0.001) by 64% compared with LL control plants (Fig. 2C). In contrast, after a single 4h HL exposure, followed by photosynthesis measurements in the next photoperiod, no increase in *A*_sat_ was observed (Fig. 2C). In a separate series of experiments, the measurement of *A* over a range of internal leaf CO_2_ concentrations (*Ci*) also showed that the maximal CO_2_-saturated rate of photosynthesis (*A*_max_) increased by 31% (P < 0.002) after 5 daily HL exposures (Fig. 2D). This confirmed that repeated HL exposures did not solely affect stomatal behaviour but brought about an increase in foliar photosynthetic capacity. The changes in CF parameters by day 5 of HL treatments observed in the previous experiments (Fig. 2A) occurred also in larger older leaves that were required for gas exchange measurements (Supplemental Figure 2C; see Methods).

In summary, increased *A*_sat_ and *A*_max_ after 5 days of repeated HL exposure (Fig. 2C, D) was accompanied by a highly significant increase in Fq’/Fm’ (Fig. 2A, Supplemental Figure 2C; P < 0.0001, ANOVA and Tukey HSD; Supplemental Data Set 3) reflecting an increased photochemical efficiency to support dynamic acclimation. Therefore, a substantial (>40%; typically using the median 800 µmol m^-2^ s^-1^ actinic PPFD value) change in Fq’/Fm’ between days 1 and 5 of repeated HL was subsequently used as a more convenient image-based measurement of the establishment of dynamic acclimation and consequent increased photosynthetic capacity.

### Dynamic statistical modelling infers a BBX32-centric network of HL-regulated transcription factor genes

The HL time series data were used to infer gene regulatory networks (GRNs) using VBSSM (Beal et al., 2005; Penfold and Wild 2011). We chose VBSSM because it has been demonstrated to infer known GRNs from temporal gene expression data and to infer novel GRNs whose highly connected genes (nodes) have subsequently been shown experimentally to have a novel and important function (Beal et al., 2005; Penfold and Wild, 2011; Breeze et al., 2011; Penfold and Buchanan-Wollaston, 2014; Windram and Denby 2015; Bechtold et al., 2016). However, due to the limited number of time points, we opted to infer networks for around 100 genes or probes in order to avoid overfitting by constraining the network size (Beal et al., 2005; Allahverdeyeva et al., 2015; Windram and Denby, 2015; Bechtold et al., 2016). To accommodate this limitation, we focussed on DEGs coding for transcription (co-) factors (TFs). We reasoned that TF GRNs would control the expression of a wide network of genes and by inferring GRNs this would allow us to identify and focus on the most connected TF genes, often termed hub genes (Windram and Denby, 2015; Albihlal et al., 2018). Consequently, we reasoned that upstream hub TF genes would directly and indirectly regulate the expression of a sufficiently large number of genes to influence whole leaf HL responses and acclimation phenotypes. Therefore, the intention was to screen highly connected candidate regulatory hub TF genes directly for their impact upon whole plant dynamic acclimation.

It was estimated that there were 371 HL DEGs coding for TFs or transcription co-factors (Supplemental Data Set 4). To narrow our selection further, comparisons were made between the 43 HL temporal clusters (Fig. 1A; Supplemental Data Set 1) and 14 publicly available transcriptomics data sets or meta-analyses of such data for treatments or mutants perturbed in chloroplast-to-nucleus and ROS-mediated signalling (Supplemental Data Set 5). On a cluster-by-cluster basis, the highest number of significant (P < 0.00001) overlaps in clusters 1, 2, 3, 5, 6, 9, 10, 14, 16, 17 and 27 were encountered with *phyA/phyB* DEGs (Supplemental Data Set 5) among which genes involved in chloroplast-to-nucleus (retrograde) signalling had been identified (Shikata et al., 2014). This observation suggested that photoreceptor-mediated regulation of HL-responsive genes was highly represented in the time series transcriptomics dataset. Therefore, we examined whether photoreceptor-regulated TF and co-transcription factor genes (Shikata et al., 2014; Dong et al., 2014) were also over-represented in the HL dataset. This was the case. Ninety-one (91) photoreceptor- and light-regulated TF DEGs were over-represented in the time series transcriptomics data, irrespective of which temporal cluster they were drawn from (P= 1.4E-06; Hypergeometric Distribution Test, Supplemental Data Set 4). The HL time series expression data from these 91 genes were used to infer networks with VBSSM.

The first inferred network for HL revealed *LATE ELONGATED HYPOCOTYL* (*LHY*) as the most highly connected gene (Supplemental Fig. 4A). However, mutant *lhy-21* plants were not perturbed in dynamic acclimation (Supplemental Fig. 4B). Therefore, the VBSSM modelling was re-iterated but omitting *LHY* expression data. This inferred a 47 node-HL network centred on *BBX32* (Fig. 3). The transcript levels of the 12 most connected nodes (≥ 3 edges) across the time series, under LL and HL conditions, is shown in Supplemental Figure 5 and shows the diversity of expression patterns derived from the temporal clusters (Supplemental Data Set 1; Fig. 1A; B).

**Figure 3.**
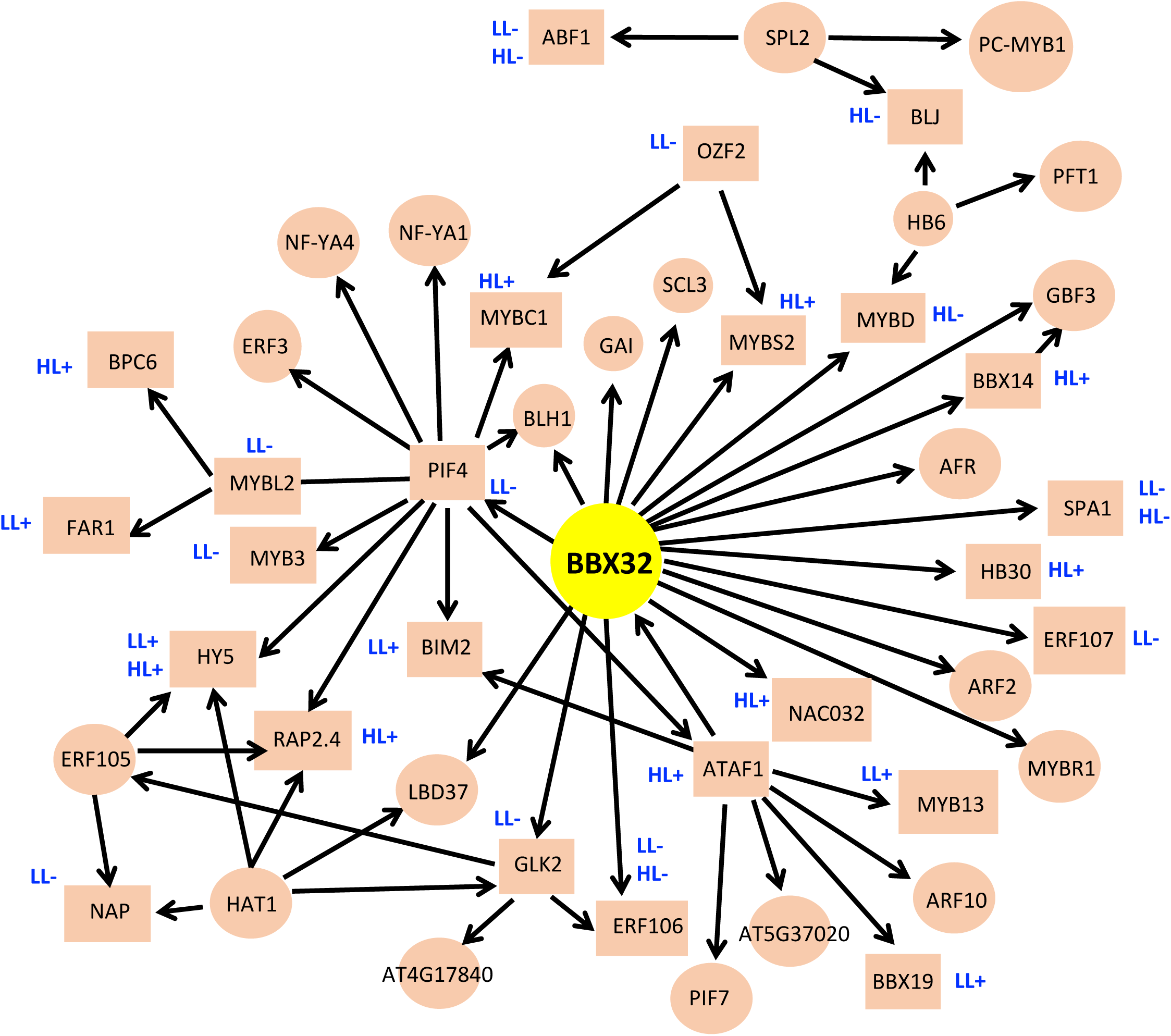
**Inferred HL gene regulatory network centered on BBX32.** The network shown was generated from the time series expression data for HL DEGs. The DEGs code for transcription (co)factors that are also light- and/or *PHYA/PHYB* regulated in de-etiolating seedlings. The network was generated using VBSSM (threshold z-score = 2.33; see Methods) and initially visualised using Cytoscape (v3.3.2; Shannon et al., 2003) but re-drawn manually to improve clarity. The network shown is from the second iteration of the modelling, which omitted expression data for *LHY* (First iteration; Supplemental Fig. 4A). The genes depicted in rectangular nodes were responsive to *BBX32* over-expression in HL and / or LL exposed leaves by showing significantly (P< 0.05; Tukey HSD) higher (+) or lower (+) transcript abundance than Col-0 (see Fig. 5). Locus codes for the network genes can be found in Methods.

### BBX32 is a negative regulator of dynamic acclimation

Dynamic acclimation was tested in two independent *BBX32* over-expressing (*BBX32*-OE) genotypes (BBX32-10 and BBX32-12) and a T-DNA insertion mutant (*bbx32-1*; see Methods and Holtan et al., 2011). *BBX32-*OE plants showed a highly significant impairment of dynamic acclimation (Fig. 4A; Supplemental Data Set 6). In contrast, *bbx32-1* plants showed a weak but significant accelerated dynamic acclimation phenotype (Fig. 4B; Supplemental Data Set 6). We define an accelerated acclimation phenotype as a significant enhancement of PSII operating efficiency over one or more days in the 5d serial HL treatment. The strong negative impact of *BBX32* over-expression on dynamic acclimation was confirmed subsequently by showing a significant inhibition of photosynthetic capacity (A_sat_) after 5 days of daily 4h HL exposure (Fig. 4C). Consequently, it was concluded that *BBX32* is a negative regulator of dynamic acclimation.

**Figure 4.**
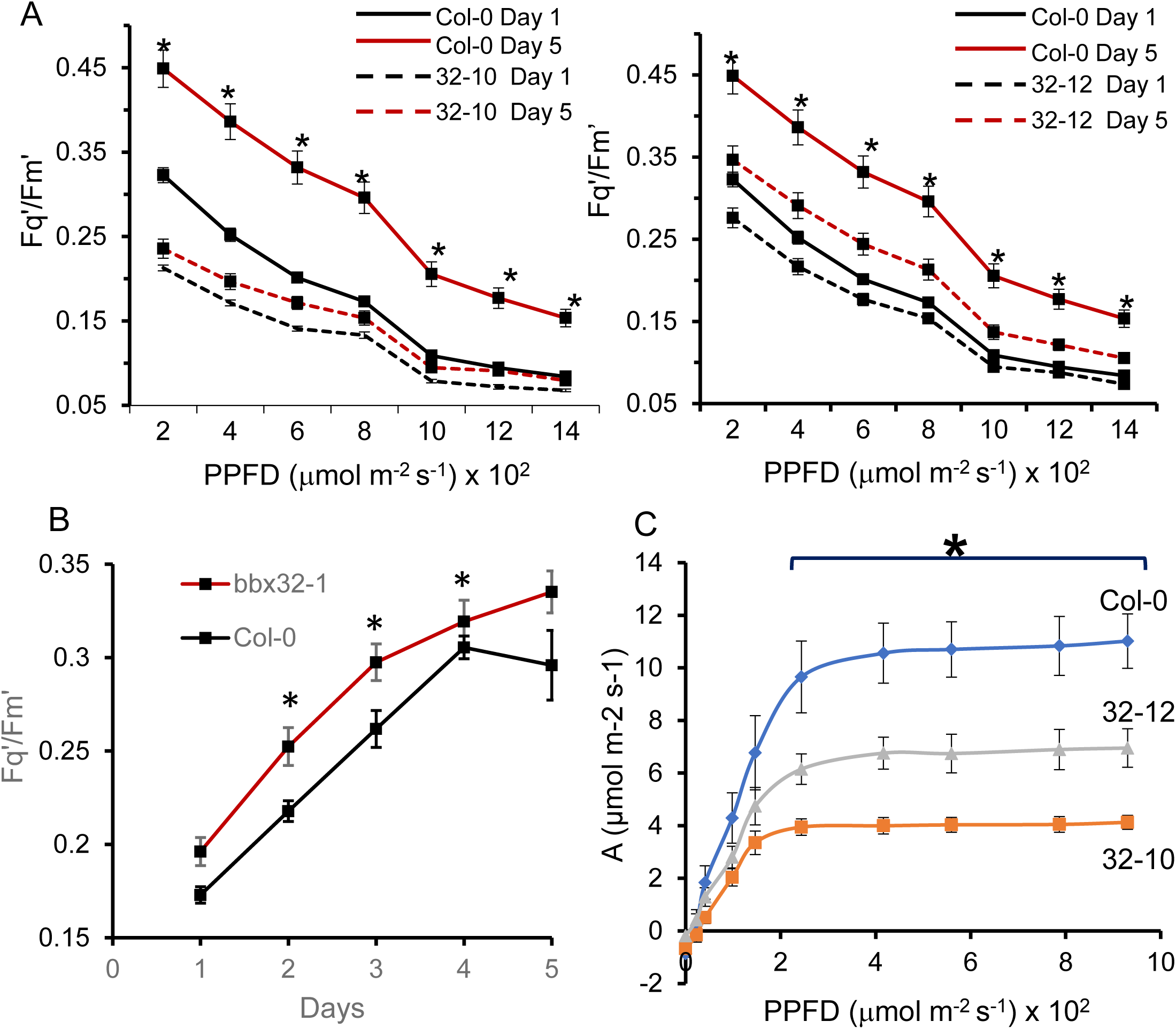
**Dynamic acclimation in *BBX32-*OE and *bbx32-1* plants.** Fq’/Fm’ values determined from images of ≥ 4 mature leaves from 8 plants (24-28 dpg) over 2 experiments (means ± SE) which had first been exposed to 4h HL each day for 5 consecutive days (see Methods and legend of Figure 2). CF parameter values were collected at a range of actinic PPFDs (as indicated) at the end of each daily HL exposure. **(A)** Fq’/Fm’ values at day 1 (black lines) and day 5 (red lines) for mutant or OE plants (dashed line) and Col-0 (solid line) of the HL treatments for BBX32-10 and BBX32-12. Asterisks indicate difference between mutant genotype and Col-0 at day 5 (P < 0.01; ANOVA and TukeyHSD). **(B)** Daily Fq’/Fm’ values at 800 µmol m^-2^ s^-1^ PPFD actinic light of *bbx32-1* compared with Col-0 showing differences that were significant (P <0.01) only between days 2 and 4. **(C)** Photosynthesis plotted as CO_2_ assimilation rate (*A*) as a function of incident PPFD in mature leaf 7 of LL-grown BBX32-10 (green line) and BBX32-12 (red line) compared to Col-0 (blue line) plants. Data are the mean ± SE; n = 4 for each genotype at 49 dpg; Asterisk indicates significant differences (P < 0.02; covariant T and two-tailed F tests) between Col-0 and BBX32-10 and BBX32-12 at a given PPFD. Leaf *A*, as a function of PPFD, was determined by Infra-Red Gas Analysis (see Methods).

#### Transcriptomics provides a partial verification of the BBX32 HL TF network

In order to explore further the connections depicted in the network model (Fig. 3), massively parallel RNA sequencing (RNAseq; Supplemental Data Set 7) was carried out (see Methods; GEO; GSE158898) to profile the foliar transcriptome of fully expanded leaves of Col-0 and *BBX32-*OE plants exposed to 3.5h HL in comparison with LL controls. From the RNAseq data, the transcript levels of 25 of 47 constituent genes in the inferred network were significantly altered by constitutive *BBX3*2 over-expression compared with Col-0 plants in LL and/or HL (Fig. 5). Therefore, these data partly validated this (co)TF GRN and suggested that *BBX32* engages in both positive and negative regulation of the other TF genes in this network (Fig. 3; Fig. 5).

**Figure 5.**
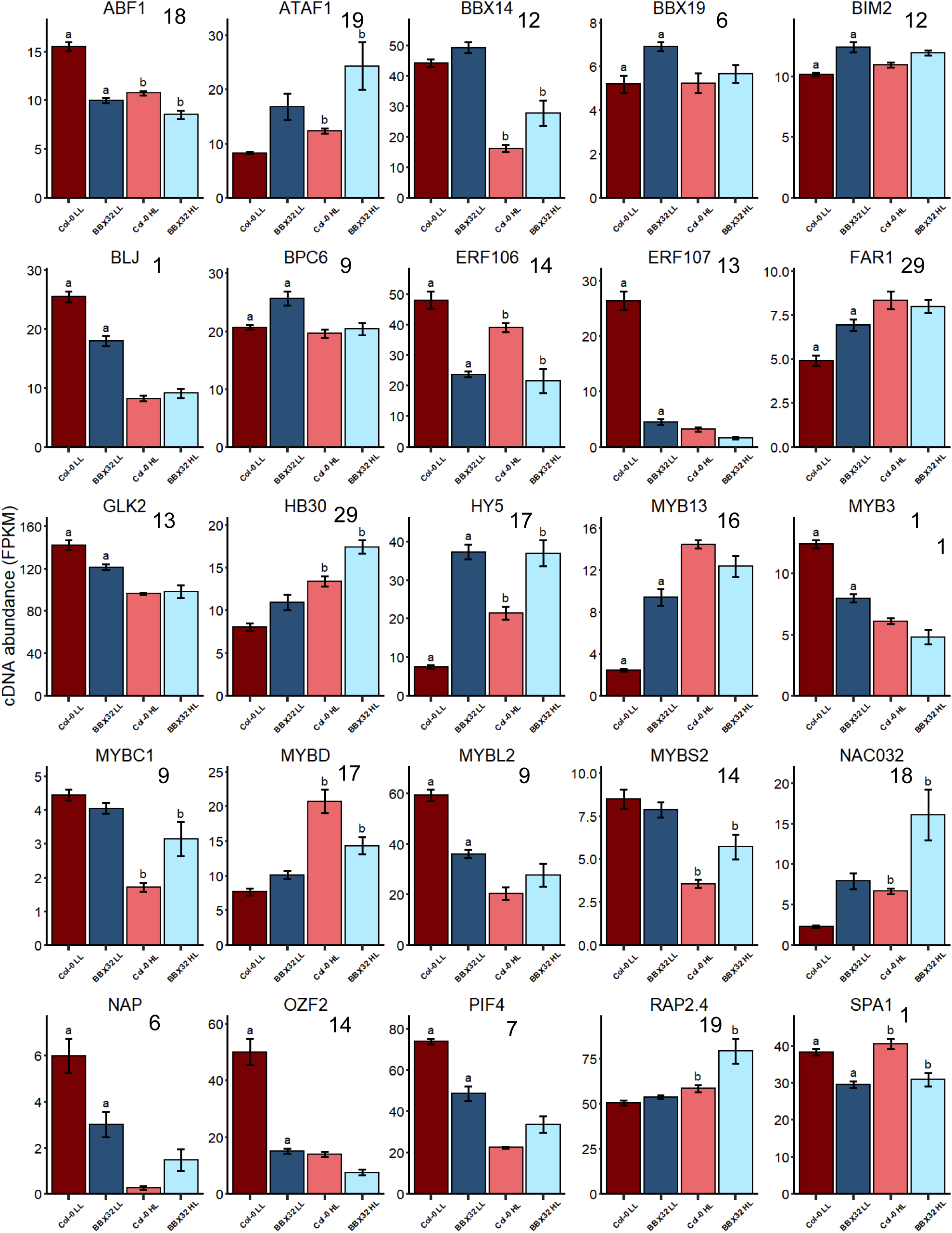
**Partial validation of the BBX32-centric inferred gene regulatory network.** The expression of 25 of the 47 TF genes in the inferred network showing the effect of *BBX32* over-expression. All the genes displayed significant differences (p <0.05; Tukey HSD) in cDNA abundance in *BBX32*-OE plants compared with Col-0 under LL (suffix “a”) and/or HL (suffix “b”) conditions. Tabulated FPKM data for these genes can be found in Supplemental Data Set 7. Color codes are dark red and blue Col-0 and *BBX32*-OE plants in LL respectively, salmon pink and light blue are Col-0 and BBX32-OE plants in HL. The cluster number for each gene is shown on each graph.

### The transcriptome of BBX32OE plants links initial responses to HL with dynamic acclimation

The greatly impaired ability of *BBX32*-OE plants to undergo dynamic acclimation (Fig. 4A, C), prompted an analysis of the RNAseq data on the impact of *BBX32* over-expression on the transcript levels of photosynthesis-associated genes (PhAGs; https://www.kegg.jp/dbget-bin/www_bget?pathway+ath00195). There was a clear influence of *BBX32* over-expression under LL and HL on the transcript levels of a range of transcripts coding for LH Antenna proteins, Calvin-Benson cycle enzymes and components of photosynthetic electron transport, PSI and PSII (Fig. 6; Supplemental Data Set 8). We concluded that these and other transcripts affected in *BBX32*-OE plants may reflect their perturbed photosynthetic physiology.

**Figure 6.**
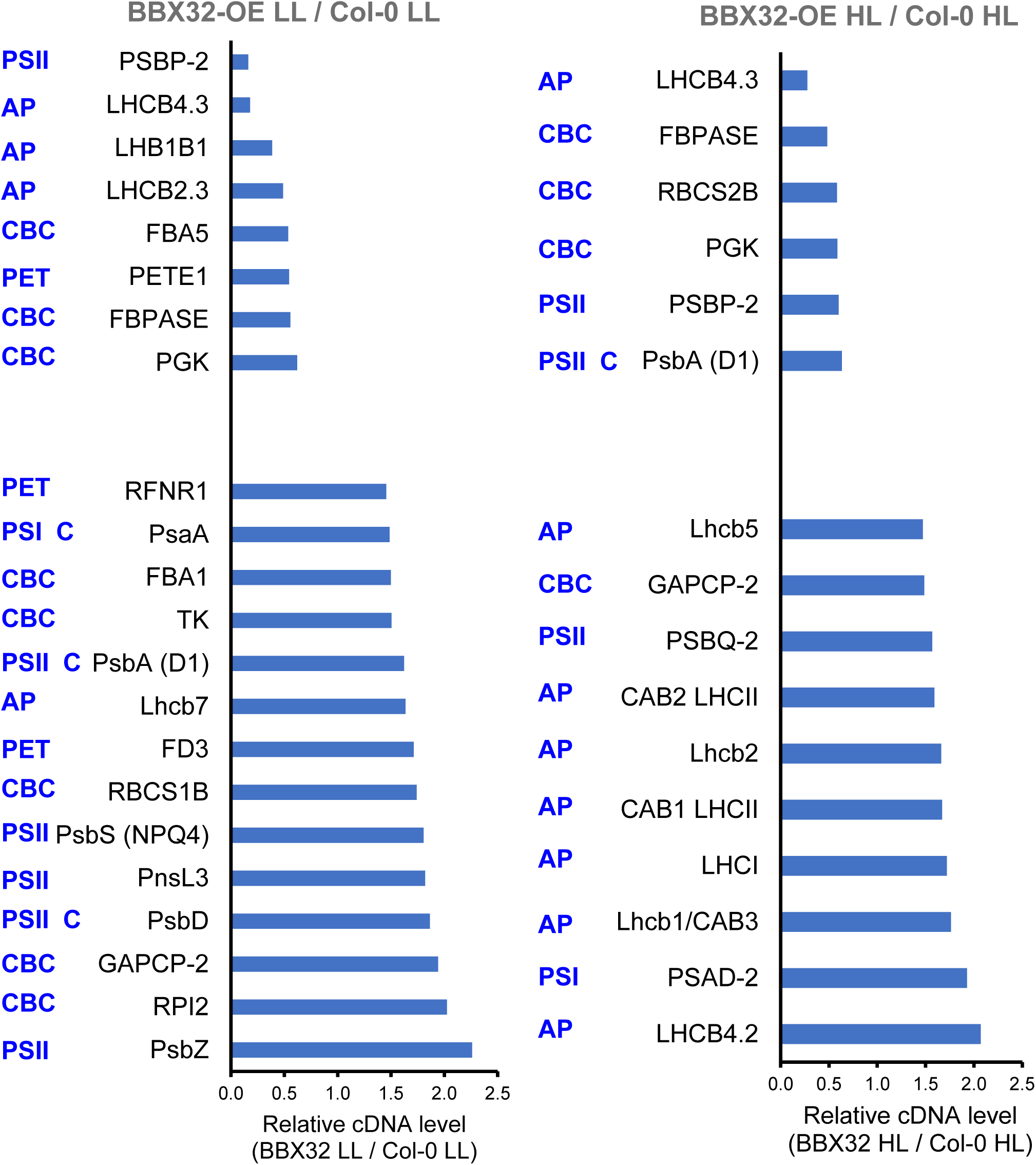
**BBX32 over-expression in LL and HL-exposed leaves perturbs transcript level of photosynthesis-associated genes.** Using RNAseq data, relative cDNA abundance of *BBX32*-OE compared with Col-0 of photosynthesis-associated genes was determined under LL and 3.5 h HL exposure. The transcripts encoding the above proteins all displayed at >1.45 -fold greater or lesser abundance in fully expanded leaves of *BBX32* - OE plants. The values are calculated from mean FPKM values (n=4) and difference between Col-0 LL and Col-0 HL were significant (P adj. < 0.05). In blue are the designated classifications for photosynthesis associated genes (https://www.kegg.jp/dbget-bin/www_bget?pathway+ath00195): AP, Antenna Protein; CBC, Calvin-Benson cycle enzyme; PET, photosynthetic electron transport protein; PSI and PSII, Photosystem I and II component proteins respectively. Most proteins are nuclear encoded but those marked with the suffix “C” are plastid encoded. Locus codes for the genes can be found in Methods.

This widespread disruption of PhAG transcript levels led us to examine the impact of *BBX32* over-expression on other cellular processes. In the RNAseq experiment, of the 2903 genes whose transcript levels were HL responsive (Padj. < 0.05; ≥ 2-fold differentially expressed; Supplemental Data Set 7), *BBX32* over-expression perturbed the transcript levels of 32% and 15% of them in LL and HL conditions respectively (Fig. 7A; Supplemental Data Set 7). The HL/LL Col-0 DEGs were enriched for 35 GO BP terms (Supplemental Data Set 9) and 26 of them were also significantly over-represented in the *BBX32*-OE / Col-0 LL and *BBX32*-OE / Col-0 HL DEGs (Supplemental Data Set 9). These shared GO groups all describe responses to various abiotic and biotic stresses or response to endogenous stimuli such as salicylic acid or H_2_O_2_. This analysis indicates that *BBX32* influences a wide range of cellular responses to stress, which includes regulation of genes associated with basal immunity to infection.

**Figure 7.**
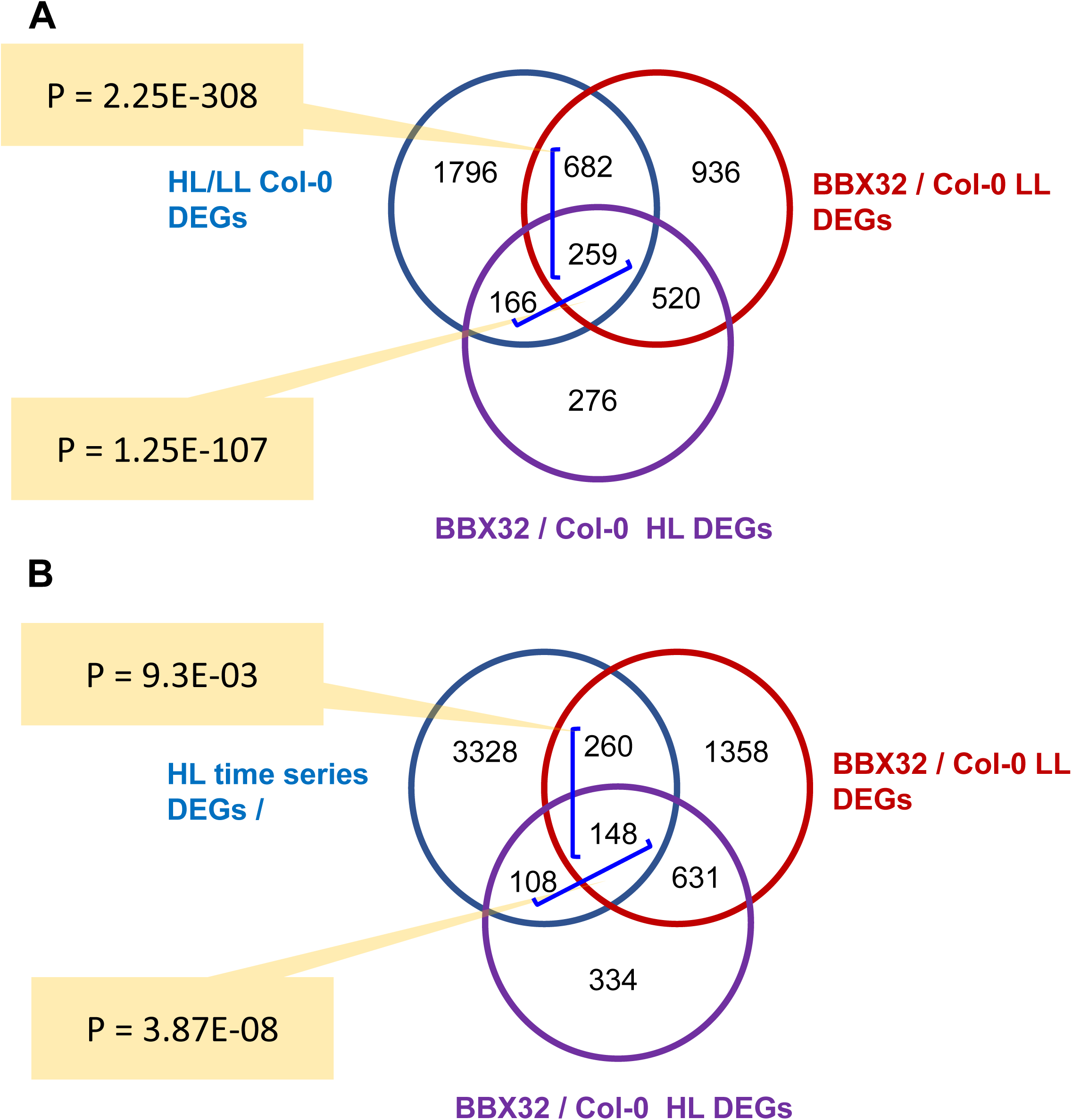
**Comparisons of genes affected by BBX32 over-expression with DEGs responsive to HL in Col-0.** **(A)** Venn diagram of overlapping DEGs between Col-0 and BBX32-OE plants under LL and HL conditions compared with DEGs responsive in Col-0 to 3.5h HL and generated by RNAseq. The relevant 3 groups of DEGs can be found in Supplemental Data Set 7. **(B)** Venn diagram as in (A) except the BBX32-OE DEGs were compared with the time series HL DEGs in Supplemental Data Set 1, which were derived from microarray-based transcriptomics data (see Results and Methods). The DEGs in the overlapping segments are listed in Supplemental Data Set 10.

The DEGs from *BBX32*-OE HL and LL treated plants were also compared with the 3844 time-series HL DEGs (Fig. 7B; Supplemental Data Set 2; Supplemental Data Set 10). Although the number of overlapping genes was lower (Fig. 7B), *BBX32*-OE HL DEGs again confirmed enrichment for a range of GO terms that describe generic responses to environmental stress (Supplemental Data Set 10). However, the 408 *BBX32*-OE LL DEGs also differentially expressed in the HL time series dataset, also revealed significant enrichment (FDR < 0.05) of a range of additional functions (Supplemental Data Set 10) including glucosinolate and glycosinolate metabolism (GO:0019760, GO:0050896, GO:006143, GO:0019757, GO:0016144, GO:0019761, GO:0019758), cell wall thickening (GO:0052543, GO:0052386) and callose deposition (GO:0052543, GO:0052545). Down-regulation of these groups of genes in the HL time series data (Supplemental Data Sets 1 and 2) may reflect a re-distribution of resources towards dynamic acclimation and away from basal immunity (see above and Discussion). The observations here suggest *BBX32* may play a regulatory role in these processes (see Discussion) but also reinforces that *BBX32* influences immediate responses before or during a single exposure to HL.

### CRY1- and HY5- regulated control of dynamic acclimation

*BBX32* has been proposed to be a negative regulator of the integration of light signals from phytochromes (PHYs) and cryptochromes (CRYs) during photomorphogenesis (Holtan et al., 2011; Gangappa and Botto, 2014). *BBX32*-OE seedlings display a long hypocotyl phenotype in the light like photoreceptor mutants and mutations in *LONG HYPOCOTYL5* (*HY5*; Holtan et al., 2011). Notably, *HY5* is a member of the BBX32-centric GRN (Fig. 3; Fig. 5) and along with *CRY1*, has also been implicated in influencing the expression of HL-inducible gene expression (Kleine et al., 2007; Shaikali et al., 2012; Chen et al., 2013). Furthermore, *PHYA*-, *PHYB*- and *CRY1*-mediated signaling was proposed to regulate maximum photosynthetic capacity in plants grown in a range of PPFDs (Walters et al 1999; see Introduction). These considerations prompted us to test dynamic acclimation in photoreceptor-defective and *hy5* mutants.

No significant impact of *PHYA* or *PHYB* on acclimation was observed (Supplemental Fig. 6A, B). In contrast, *cry1* mutants almost completely failed to undergo any dynamic acclimation (Fig. 8A, B), whereas *cry2-1* was not impaired (Supplemental Fig. 6C). One of the *cry1* mutants shown (*cry1*M32; Fig. 8B) arose serendipitously from a screening of T-DNA insertion mutants in genes coding for 7-transmembrane proteins that had been postulated to be implicated in HL-mediated G protein signaling (Galvez-Valdivieso et al., 2009; Gorecka et al., 2014). However, the one mutant recovered from this screening, was shown subsequently to be deficient in dynamic acclimation due to a disabling second site mutation in *CRY1* (see Methods). Since the defective acclimation phenotype was identified prior to knowing the identity of the causal mutation we took this to be forward genetic evidence of the importance of *CRY1* in dynamic acclimation in mature leaves.

**Figure 8.**
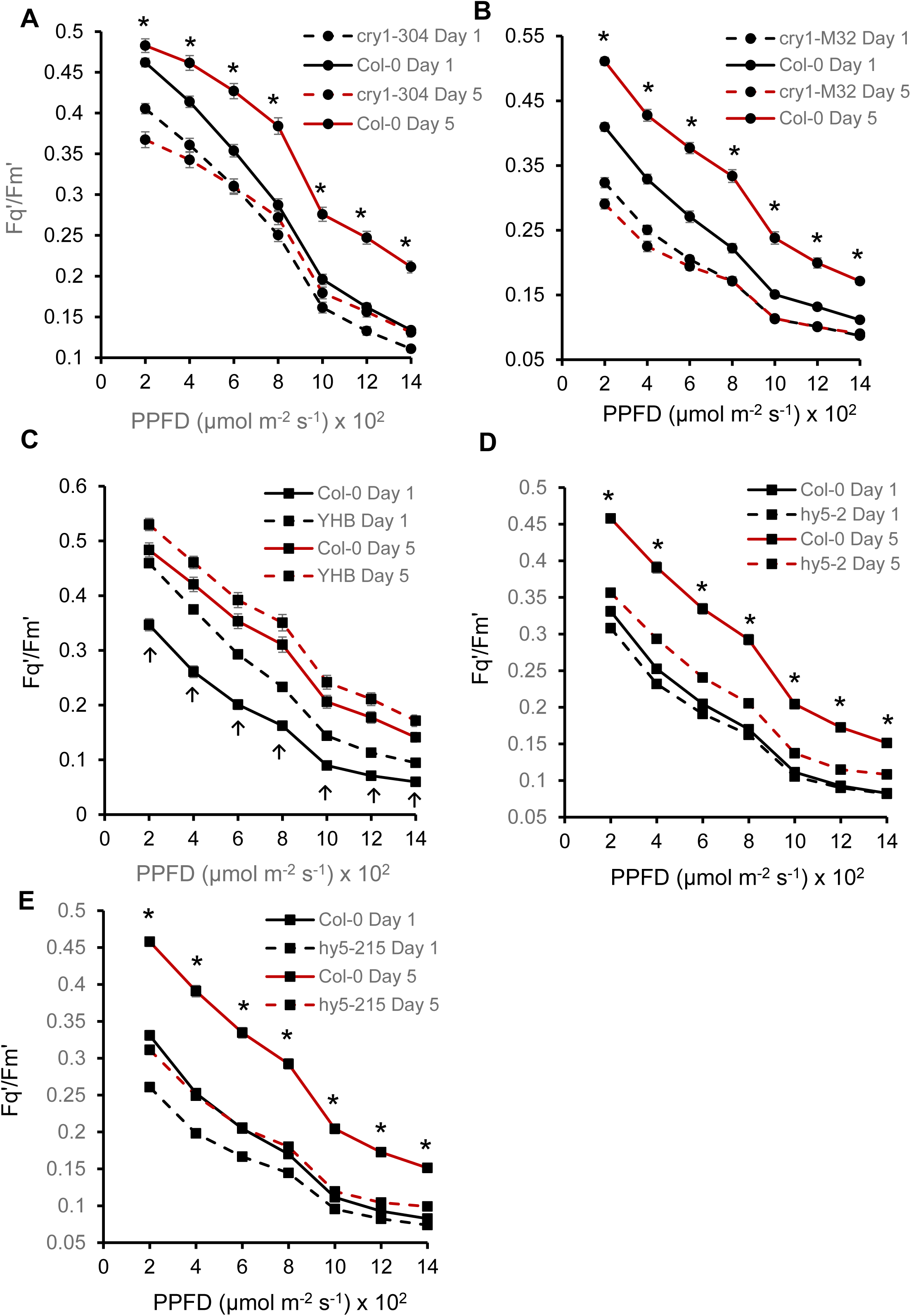
**Dynamic acclimation of photoreceptor and *HY5* mutants.** The plots show the PSII operating efficiencies (Fq’/Fm’) determined from CF images of ≥ 4 mature leaves from 8 plants (24-28 dpg) over 2 experiments (means ± SE). The plants had been exposed to 4h HL each day for 5 consecutive days (see Methods and legend of Figure 2). CF parameter values were collected at a range of actinic PPFDs (as indicated on the x-axis) at the end of days 1 and 5 of HL. The Fq’/Fm’ values at day 1 (black lines) and day 5 (red lines) for mutant plants (dashed line) and Col-0 (solid line) of the HL treatments for (A) *cry1-304*, (B) *cry1-M32*, (C) *YHB*, (D) *hy5-2* and (E) *hy5-215.* Asterisks (panels A, B, D, E) indicate significant difference between mutant compared with Col-0 at day 5 (P < 0.01; ANOVA and TukeyHSD). Upward arrows (panel C) indicate significant difference between YHB and Col-0 at day 1 (P < 0.01; ANOVA and TukeyHSD).

The light environment used to grow plants for this study and subject to HL was enriched for blue wavelengths (Supplemental Figure 7; see Discussion). Therefore, we considered the possibility that a role for phytochromes in dynamic acclimation could be obscured, favouring a predominance of CRY1 under our growth conditions. To test this notion, a mutant harbouring a constitutively active form of PHYB, phyBY276H (YHB) in a Col-0 background (Jones et al., 2015) was tested for dynamic acclimation (Fig. 8C). This mutant exhibited a higher PSII operating efficiency than Col-0 after 1 day of HL exposure. This accelerated acclimation phenotype is in keeping with being a constitutively active positive regulator of dynamic acclimation.

Mutants defective in *HY5* function were strongly impaired in dynamic acclimation (Fig. 8D, E) consistent with being a member both of a BBX32-centric GRN and being a positive regulator of CRY1-mediated dynamic acclimation (Fig 8A, B).

### *COP1, PIF* and *SPA* genes regulate dynamic acclimation

In both photomorphogenesis and shade avoidance responses, the transduction of signals from photoreceptors is mediated *via* one or more DET/COP/FUS regulatory complexes (Lau and Deng, 2012), which act as platforms for the post-translational control of the levels of HY5 and the integration into the signaling of the TFs PHYTOCHROME INTERACTING FACTORS (PIFs) and regulatory proteins SUPPRESSOR OF PHYA-105 (SPA) (Hardtke et al 2000; Toledo-Ortiz et al 2003; Lian et al 2011; Dong et al 2014; Huang et al 2014; Lau and Deng 2012; Gangappa and Botto, 2016; Hoecker 2017; Pham et al 2018; Lau et al., 2019). CRY1 and PIFs have been shown also to physically interact independent of COP/DET/FUS (Pedmale et al., 2016; Ma et al., 2016). In the VBSSM-inferred GRN, PIF4 and SPA1 were predicted to have a regulatory connection to BBX32 (Fig. 3; Fig.5). Significantly, 187 HL time series DEGs overlapped (P = 0.0018; Hypergeometric Distribution Test) with a set of 1120 genes identified as commonly regulated by *SPA1, 2, 3, 4, PIF1, 3, 4, 5* and *COP1* in de-etiolating and light-exposed seedlings (Pham et al., 2018). Interestingly, the most significant GO Biological Process function coded by these overlapping genes was Photosynthesis (GO: 0015979; FDR = 2.7 E-17; Supplemental Data Set 11).

*Cop1-4* plants, despite a severely dwarfed shoot morphology (Fig. 9A; Deng and Quail, 1992; Gangappa and Kumar, 2018), displayed a highly elevated PSII operating efficiency (Fq’/Fm’) by day 1 of the HL acclimation regime compared with Col-0 (Fig. 9B) like the HL response of YHB plants (Fig. 8C). In contrast, despite a similar dwarf shoot morphology (Fig. 9A), *det1-1* displayed no defect in dynamic acclimation (Fig. 9D). This strongly suggests that the dynamic acclimation response of chloroplasts is independent of shoot size and that these two traits are not coupled. Furthermore, *spa1/spa2/spa3 (spa1,2,3*) plants also displayed an accelerated acclimation phenotype (Fig. 9C). Therefore, it was concluded that one or more type of COP1/SPA complex (Huang et al., 2014; Hoecker, 2017) are negative regulators of dynamic acclimation and that *DET1* plays no role in dynamic acclimation.

**Figure 9.**
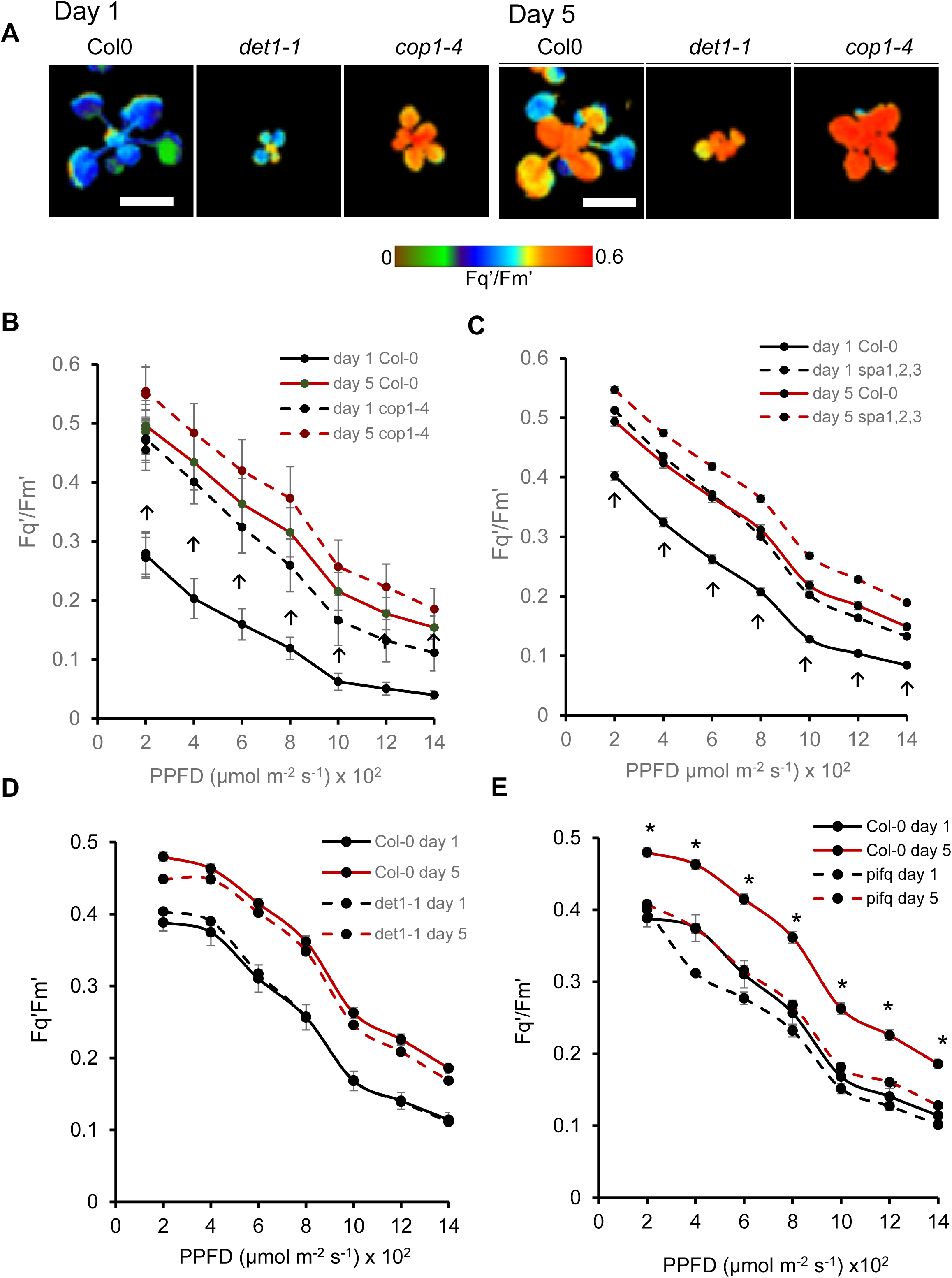
**Dynamic acclimation of photoreceptor signal transduction mutants.** **(A)** Photosynthetic efficiency of the same single representative Col-0, *cop1-4* and *det1-1* plants after 1 and 5 days of daily 4h HL exposure. The CF images are of Fq’/Fm’ (PSII operating efficiency) at a 400 µmol m^-2^ s^-1^ actinic PPFD. (B – E) The plots show the PSII operating efficiencies (Fq’/Fm’) determined from CF images of from 8 plants (24-28 dpg) over 2 experiments (means ± SE). The plants had been exposed to 4h HL each day for 5 consecutive days (see Methods and legend of Figure 2). Note that because of the size of the *cop1-4*, *pifq* and *det1-1* plants, data were collected from whole rosettes rather than from mature leaves. CF parameter values were collected at a range of actinic PPFDs (as indicated on the x-axis) at the end of days 1 and 5 of HL. The Fq’/Fm’ values at day 1 (black lines) and day 5 (red lines) for mutant plants (dashed line) and Col-0 (solid line) of the HL treatments for (B) *cop1-4*, (C) *spa1,2,3*, (D) *det1-1* and (E) *pifQ.* Asterisks (panel E) indicate significant difference between mutant compared with Col-0 at day 5 (P < 0.01; ANOVA and TukeyHSD). Upward arrows (panels B, C) indicate significant difference between mutants and Col-0 at day 1 (P < 0.01; ANOVA and TukeyHSD).

There is a high degree of redundancy among the PIF TF family and therefore a quadruple null mutant of *PIF1*,*3*, *4* and *5* (hereafter called *pifq*; Leivar et al., 2008) was tested for its capacity for dynamic acclimation. These plants displayed a severe dwarf phenotype as previously described (Leivar et al., 2008), but also a significant inhibition of dynamic acclimation (Fig. 9E). In contrast, the dynamic acclimation of a single mutant allele of *PIF4* (*pif4-2*) was normal (Supplemental Fig. 6D).

## DISCUSSION

### The time series HL transcriptomics data indicates the initiation of dynamic acclimation processes

The exposure to a 7.5-fold increase in PPFD (HL; see Methods) presents both a threat and an opportunity to the plants in this study. The threat comes from the possibility that the PPFD will continue to increase and render the plant susceptible to irreversible photoinhibition. The opportunity comes from enhancing photosynthesis by initiating dynamic acclimation (Fig. 2A-D). Accompanying enhanced photosynthesis was also a lowering of reliance on the dissipation of excitation energy using NPQ (Supplemental Fig. 2A), which can limit plant productivity (Kromdiijk et al., 2016).

The adaptation to a potential increase in photooxidative stress and photoinhibition (see Introduction) is the early (≤ 1h) strong but transient change in transcript abundance of 257 genes in clusters 21-26, upon exposure to HL. Clusters 22, 23, 25 and 26 include among them 64 known genes that promote abiotic stress tolerance (Fig. 1A, B; cluster 23 in Supplemental Fig. 1; Supplemental Fig. 2C; Supplemental Data Set 1; Supplemental Data Set 2). The transiently enhanced expression of these genes presumably allows the plant to overcome any potential initial detrimental effects of the HL exposure, as many other studies have reported (e.g. Ball et al., 2004; Gadjev et al., 2006; Ramel et al., 2012; 2013; Willems et al., 2016; Crisp et al., 2017; Huang et al., 2019).

Coordinated alteration in specific biological processes was evident in some clusters. Down-regulated clusters include those collectively associated with aspects of basal or innate resistance to pathogens (Underwood, 2012; Piasecka et al., 2015). Examples include genes coding for cell wall modifications and callose deposition (cluster 1), defense response to bacteria (cluster 3) and glucosinolate / glycosinolate biosynthesis (cluster 10). In this study, plants were grown at a PPFD below their light saturated rate of photosynthesis (Asat; Fig. 2C; see Methods). Plants grown under such light-limiting conditions may initially have to re-allocate resources away from some cellular processes in order to begin acclimation and take advantage of a sustained increase in PPFD. Photosynthetically active expanded but not senescing leaves, such as leaf 7 used here (see Methods), have been suggested to maintain a higher degree of poising of immunity to respond to biotic stress compared with abiotic stress (Berens et al., 2019). Therefore, in a converse situation where a potential abiotic stress threat emerges, the diversion of resources from defense against pathogens may be an appropriate response. Meanwhile, among the DEG time series clusters whose transcript levels increased at various points in the experiment, are those that could be preparing the leaf to increase its photosynthetic and metabolic capacity in order to begin acclimation (Eberhard et al., 2008; Dietz, 2015). Genes in over-represented GO BP terms included those involved in macromolecule synthesis and especially translation (clusters 41-43) and related metabolic processes such as enhanced amino acid and organic acid biosynthesis (cluster 39).

A single exposure to 4h HL is not sufficient to induce dynamic acclimation at the physiological level, requiring, under our conditions, a further 3 daily episodes of 4h HL for this to begin to occur (Fig. 2A-D). Our experience is consistent with a previous study where dynamic acclimation took around 5 days to be fully manifested and 2-3 days to discern any change in photosynthesis rates after a permanent shift from a photoperiod PPFD of 100 µmol m^-2^ s^-1^ to 400 µmol m^-2^ s^-1^ (Athanasiou et al., 2010). However, it should be noted that the HL regime used did not produce dynamic acclimation for the Col-0 accession but did for others such as Ws-2 and Ler-0 (Athanasiou et al., 2010). In contrast, the shorter more intense PPFDs used in this study induced dynamic acclimation in Col-0 and also Ws-0 (Fig. 2A-D; Fig. 4A-D; Fig. 8A-E; Supplemental Fig. 4B).

### *BBX32* connects a range of cellular processes to dynamic acclimation

Of all the comparisons carried out with relevant transcriptomics data sets, the most extensive overlap with time series HL DEGs was with those from dark-germinated *phyA/phyB* seedlings exposed to red light (Supplemental Data Set 5; Shikata et al., 2014). While this was initially surprising because of the very different experimental conditions, earlier studies had shown a strong influence of photoreceptor genes (*CRY*s and *PHY*s) on photosynthetic capacity in Arabidopsis grown at a range of PPFDs (Walters et al., 1999) and an impact on the induction of some HL-responsive genes (Kleine et al., 2007; Shaikali et al., 2012; Guo et al., 2017). Our own and the published data above, prompted a selection of 91 light- and PHYA/B-regulated (co)TF genes (Supplemental Data Set 5). The HL time series expression data from these genes was subjected to VBSSM, which after two iterations, inferred a highly interconnected *BBX32*-centric (co)TF GRN (Fig. 3 and see Results). In the GRN, >50% of the nodes (genes) were subsequently confirmed by RNAseq to be influenced significantly in their expression by *BBX32* (Fig.3; Fig. 5).

Over-expression of *BBX32* clearly demonstrated that this gene is a negative regulator of dynamic acclimation (Fig. 4A-D) and has an extensive influence on immediate responses to HL that include processes associated with photosynthesis (Fig. 6; Supplemental Data Set 8). Notably, depressed levels of *LHCB4.3* transcript in *BBX32*-OE HL and LL plants (Fig. 6) could be important since levels of this antenna protein are closely linked to the degree of long-term acclimation to HL (Albanese et al., 2016).

*BBX32* overexpression also impacts on a range of cellular processes that can be associated with basal immunity, including multiple GO designations for glucosinolate/glycosinolate metabolism, callose deposition, responses to chitin and to pathogens (Supplemental Data Sets 7, 9 and 10). This observation is consistent with the enrichment of the same processes noted in down-regulated temporal clusters (see above; Supplemental Data Set 2) and supports our suggestion that in wild type plants, down-regulation of basal immunity may be a necessary prerequisite for successful dynamic acclimation (see above). We propose that *BBX32* control of aspects of basal immunity is part of its regulation of the initiation of dynamic acclimation.

*BBX32* showed a greater transcript abundance over LL controls at any point onwards from the 2h HL timepoint. Nevertheless, its transcript abundance was on a downward trend through the diel, paralleling its LL pattern of expression (Supplemental Fig. 5). Interestingly, while *BBX32*-OE plants displayed a 66-fold elevated *BBX32* transcript level in LL, this value reduced to 33-fold after 3.5h HL (Supplemental Data Set 7). The enhanced *BBX32* expression in these plants is driven by the CaMV 35S promoter (Holtan et al., 2011), therefore the decline in transcript abundance over a diel could indicate that a temporal post-transcriptional control operates to determine *BBX32* transcript levels.

The strong negative impact of *BBX32* over-expression upon dynamic acclimation suggested that a defective gene ought to confer a converse elevated phenotype. The mutant *bbx32-1* (see Results; Holtan et al., 2011), displayed a weakly significant trend of enhanced PSII operating efficiency compared with Col-0 between days 2 and 4 of the 5 days of 4h HL exposure (Fig. 4B). This genotype, however, is unlikely to be a null mutant. The mutagenic T-DNA is inserted such that the first 172 amino acid residues of BBX32 would still be produced and a truncated transcript spanning this region has been detected in *bbx32-1* seedlings (Holtan et al., 2011). The retained N-terminal region coded by this allele harbors the single B-Box zinc finger domain of BBX32 (Gangappa and Botto, 2014) and downstream sequences to residue 88, capable of binding at least one target protein, the transcription regulator EMBRYONIC FLOWER1 (EMF1; Park et al., 2011). The possibility of a partially functional truncated *BBX32* may explain the weak phenotype of *bbx32-1* with respect to this acclimation phenotype (Fig. 4B) and also its mild constitutive photomorphogenic phenotype in seedlings (Holtan et al., 2011).

### Two levels of control of dynamic acclimation

The time series data and the VBSSM modelling led us to identify *BBX32* and *HY5* as strong negative and positive regulators respectively of dynamic acclimation in mature leaves (Fig. 4A-D; Fig. 8D, E) and places the start of the process right at the first hours of exposure to HL. This represents new functions for these two important genes and extends their role to cover a further dimension in the interaction of the mature plant with its light environment. In seedlings, *HY5* controls the positive regulation of chlorophyll content, transcript levels of PhaGs in cool temperatures (Toledo-Ortiz et al., 2014) and the control of chloroplast development during photomorphogenesis (Ruckle et al., 2007), which suggests, along with data shown here (Fig. 6; Supplemental Data Set 8), that control of these photosynthesis-associated processes by a *BBX32/HY5*-regulatory module is retained throughout the life of the plant. Furthermore, we have established that *BBX32* fulfills the criterion of regulating both immediate responses to HL and the resulting dynamic acclimation, thus providing a link between these temporally distinct processes and experimental support for the hypothesis proposed here and by Eberhard et al. (2008; see Introduction).

The comparison drawn between the control of photomorphogenesis and dynamic acclimation was extended by establishing that *CRY1* (and possibly *PHYB*) and one or more members of the *PIF* family are positive regulators of dynamic acclimation (Fig. 8A, C; Fig. 9E), while *COP1* and one or more of *SPA1*, *SPA2* and *SPA3,* are negative regulators (Fig. 9B, C). Again, by analogy with seedling photomorphogenesis, we suggest that these genes act together to suppress dynamic acclimation under LL conditions by enabling the ubiquitin-mediated degradation of HY5 and other TFs. In HL, this suppression would be reversed by CRY1 physically interacting with and inhibiting the action of COP1/SPA protein. (Laubinger et al., 2004; Lian et al., 2011, Lau and Deng 2012; Huang et al., 2014; Gangappa and Botto, 2016; Hoecker, 2017; Pham et al., 2018; Lau et al., 2019). Consequently, CRY1 would cause the re-direction of HY5 to the activation of dynamic acclimation. However, a further adaptation may be required to slow or accelerate dynamic acclimation. For example, to fine tune the establishment of dynamic acclimation in a fluctuating light environment. We suggest under HL, when *HY5* is free of negative regulation by *COP1/SPA*, that *BBX32* is the important additional moderator of the establishment of dynamic acclimation. We speculate in the scheme in Figure 10 how this system may work and provides a basis for further studies. The transcriptional control of *HY5* and by extension, other members of the *BBX32*-centric GRN (Fig. 3; Fig. 10), could be subject to regulation by additional intracellular signals in HL, such as those from chloroplasts and hormones, serving to coordinate a range of cellular processes for dynamic acclimation (Hardkte et al, 2000; Galvez-Valdivieso et al 2009; Estavillo et al., 2011; Ramel et al., 2012; 2013; Dietz, 2015; Gangappa and Botto, 2016; Guo et al., 2016; Exposito-Rodriguez et al., 2017).

**Figure 10.**
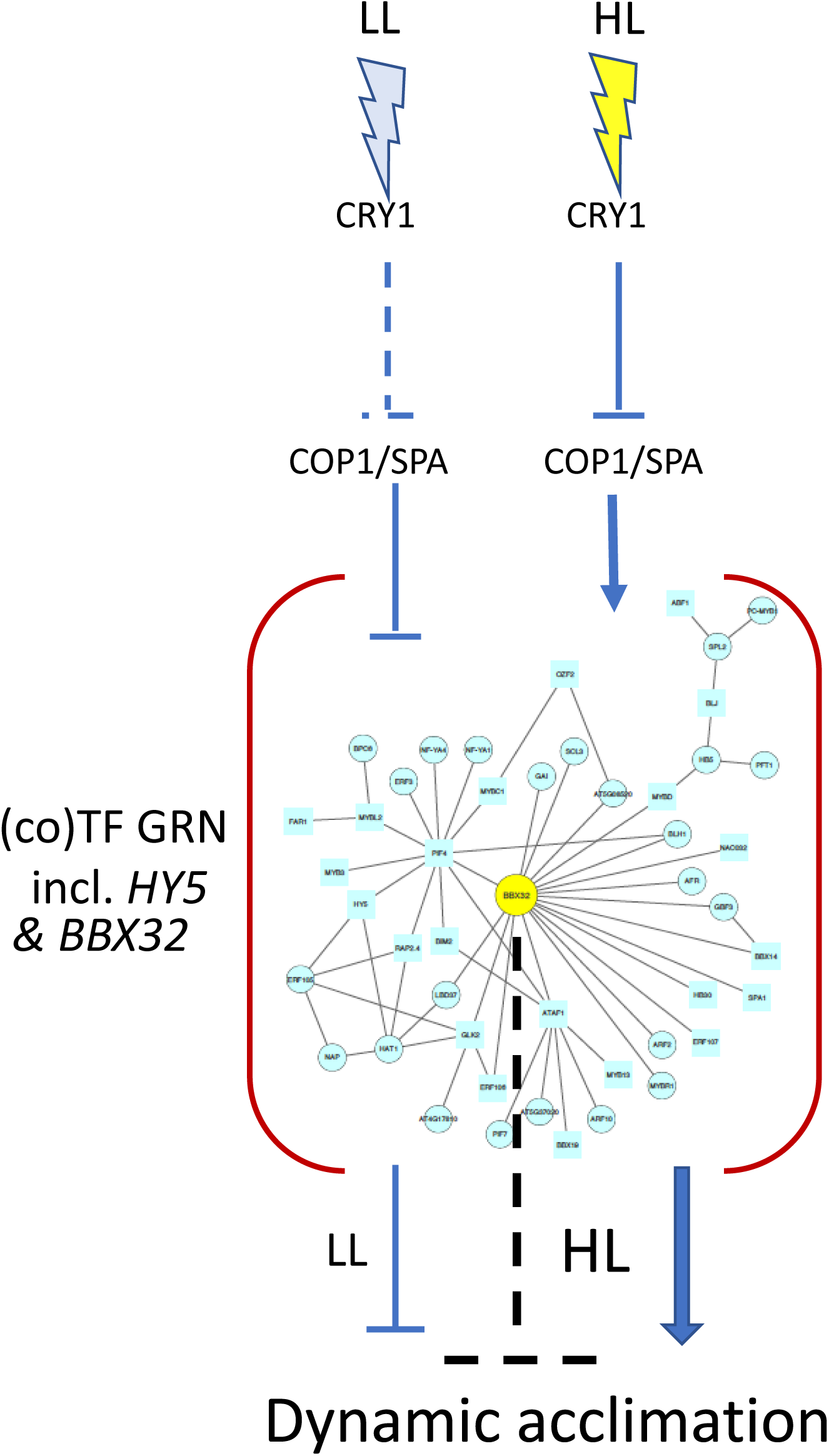
**Proposed regulation of dynamic acclimation by a B*BX32-*centric GRN.** The above scheme, while was based on this study, incorporates features from schemes published on the photoreceptor-directed control of seedling photomorphogenesis (e.g. Holtan et al., 2011; Hoecker, 2017). Under LL growth conditions to which the plant is acclimated, all or a proportion of cellular CRY1 is not active and consequently COP1/SPA acts to negatively regulate GRN members including HY5 and BBX32. Upon exposure to HL, CRY1 is activated and blocks COP1/SPA, which in turn releases the GRN ultimately leading to the establishment of dynamic acclimation. However, a degree of negative regulation of dynamic acclimation is retained (dotted inverted T) under HL conditions to allow for flexibility in potential fluctuations in the light environment.

The opposing regulation of dynamic acclimation by *BBX32* and *HY5* could mean that some form of genetic interaction between these genes drives its establishment in a manner similar to their negative and positive regulation respectively of photomorphogenesis (Datta et al., 2007; Holtan et al., 2011; Xu et al., 2014; Gangappa and Botto, 2016). However, BBX32 does not bind DNA and has been proposed to act as transcription co-factor in complexes with several TFs (Park et al., 2011; Holtan et al., 2011; Ganagappa and Botto, 2016; Tripathi et al., 2017). Of relevance here, in a tripartite complex with BBX21, BBX32 has been suggested to diminish the binding of HY5 to its target promoters (Datta et al., 2007; Holtan et al., 2011; Xu et al., 2014; Gangappa and Botto, 2016). Therefore, alongside transcriptional control of *HY5* by *BBX32*, there may also be this post-translational control of HY5 action by BBX32 during dynamic acclimation.

The proposed need for both a *CRY1/COP1/SPA-* and a *BBX32*-mediated control of dynamic acclimation (Fig.10) comes also from considerations about light quality and intensity. First, the fluence of blue light in the HL exposure used in this study would exceed the saturation of CRY1 activation, which is *ca*. 32 - 40 µmol m^-2^ s^-1^ blue light (Hoang et al., 2008; Liu et al., 2020). Therefore, while CRY1 signaling would need to be activated (i.e. on) for dynamic acclimation to happen, further signaling input may be required from other sources *via BBX32* and its GRN to modulate the degree of response. A second factor is that at high fluence, CRY1 may produce H_2_O_2_ in the nucleus (Consentino et al., 2015). H_2_O_2_ for HL signaling is primarily synthesized and exported from chloroplasts and is dependent upon an active photosynthetic electron transport chain (Exposito-Rodriguez et al., 2017; Mullineaux et al., 2018). However, this does not exclude the possibility that the HL-dependent accumulation of H_2_O_2_ in nuclei may be augmented from other sources such as photo-saturated CRY1, signals from which could be fed into the *BBX32*-centric GRN.

In contrast to Arabidopsis grown at differing PPFDs but using similar fluorescent lighting to this study (Walters et al., 1999; see Methods), no influence of *PHYA* or *PHYB* was observed on dynamic acclimation (Supplemental Fig. 6A, B). This could have been a consequence of the degree of blue light used in both the growth conditions and in applying a HL exposure (9% and 58% of total PPFD respectively; Supplemental Fig. 7; see Methods). This range of wavelengths in artificial lighting is typical of many controlled environment conditions (Naznin et al., 2019) and may have favored a response mediated by *CRY1*. The observation that plants harboring a constitutively active PHYB (YHB) allele displayed a partially accelerated acclimation phenotype (Fig. 8C) means that phytochromes could also control dynamic acclimation under some light environments and modify or interact with a *CRY1*-dependent signaling pathway (Ahmad et al., 2002; Yu et al., 2010).

A further explanation could be that the PHY mutants were altered in leaf development such that this impacted on their photosynthetic properties. Equally, we cannot rule out effects of a similar nature on *BBX32*-OE, *hy5*, *cry1*, *cop1* and *spa1,2 3* plants, but the clear lack of influence of a more severe dwarf shoot morphology on chloroplast level acclimation in *det1-1* plants argues against this (Fig. 9A, D).

### BBX32 over-expressed in Arabidopsis and soybean – control of a balance between photosynthetic capacity and leaf longevity?

Arabidopsis (A*t*)*BBX32* has been over-expressed in transgenic soybean (*Glycine max*. Merr., here called *BBX32*OE-soya) and year-on-year at different field sites produced up to an 18.5% increase in yield (Preuss et al., 2012). *BBX32*-OE Arabidopsis plants showed impaired dynamic acclimation and a consequent strong depression in Asat (Fig. 4C). This would appear to be at odds with the likelihood of an increased seed yield in the field. However, at the whole plant level, this effect may not be negative when considered as follows: *BBX32*-OE Arabidopsis plants show delayed flowering caused by the interaction of BBX32 with BBX4 (COL3), EMF1 and possibly other regulators of flowering time (Park et al., 2011; Tripathi et al., 2017). One consequence of delayed flowering is often the retardation of leaf senescence (Gan and Amasino, 1995; Wingler et al., 2010). Similarly, the *BBX32*OE-soya displayed an extended period of reproductive development associated with delayed leaf senescence thus contributing to the increased yield phenotype, similar to some “stay green” crop genotypes (Preuss et al., 2012; Kamal et al., 2019). While leaf senescence has not been measured in the *BBX32*-OE plants, nodes on the *BBX32*- centric GRN include *GOLDEN-LIKE2* (*GLK2*) and *ACTIVATING FACTOR1* (*ATAF1*; also called *NO APICAL MERISTEM ATAF1/2 CUP SHAPED COTYLEDON (CUC)2* (*NAC2*); Fig. 3; Fig. 5; Supplemental Dataset 8). These genes are implicated in the maintenance of chloroplast integrity and are determinants of the entry of Arabidopsis leaves into senescence (Waters et al., 2009; Garapati et al., 2015; Song et al., 2018).

The potential negative effect of At*BBX32* over-expression in soybean depressing maximal photosynthetic capacity may not have proved detrimental because of the way the crop is grown commercially. Modern soybean varieties are grown as a row crop to achieve a high canopy coverage that maximises the absorption of light (Shepherd et al., 2018; Koester et al., 2014). Within the canopy, the photosynthetic rate of leaves below Asat may not have been significantly affected by *AtBBX32* over-expression in soya, as observed in Arabidopsis *BBX32*-OE plants (Fig. 4D).

Furthermore, we speculate that the photosynthetic capability of leaves exposed to full sun in *BBX32*OE-soya plants, while perhaps being unable to achieve a maximal Asat, benefitted from an enhanced leaf longevity and chloroplast integrity. A further effect could also have been that like their Arabidopsis counterparts, the *BBX32*OE- soya plants had higher NPQ (Supplemental Data Set 6) perhaps linked to enhanced *PsbS* transcript levels (Fig. 6), *PsbS* being a key determinant of the qE component of NPQ (Li et al., 2000). Consequently, the *BBX32*OE-soya plants may have been less susceptible to photooxidative stress (see Introduction). Thus, while raised NPQ could have lowered photosynthetic efficiency and diverted excitation energy away from photosynthesis, this may have had a protective effect under certain field and agronomic conditions and consequently improving overall crop performance.

In summary, that a network of TF genes could control dynamic acclimation, encompassing a wide range of cellular processes, implies a complex and extensive regulation that would provide resilience and flexibility in being able to accommodate input from further intracellular and extracellular signaling. At the whole plant level, this would allow for the degree of photosynthetic capacity and acclimation in individual leaves to be adjusted according to their specific micro-environments making this acclimation a truly dynamic process.

## METHODS

### Growth conditions

Plants were grown in an 8 h photoperiod (short day) at a PPFD of 150 (± 10) µmol m^-2^ s^-1^ under fluorescent tubes (Phillips TLD 58W, 830 (warm whites); the spectrum of the light source is shown in Supplemental Figure 7), 22 ± 1 °C, 1 KPa vapour pressure deficit (VPD) and cultivation conditions as described previously (Bechtold et al., 2016; Windram et al., 2012). Unless stated otherwise, all plants were used from 35 to 40 days post-germination (dpg).

### Arabidopsis genotypes

The following Arabidopsis mutants and transgenic lines, all in a Col-0 background, have been described previously: BBX32-10, BBX32-12, *bbx32*-1 (Holtan et al., 2011), *hy5*-215 (Oyama et al., 1997), *hy5*-2 (Ruckle et al., 2007), *pifq* (Leivar et al., 2008), *cop1-4* (Deng and Quail, 1992), *det1-1* (Chory et al., 1989), *spa1/spa2/spa3* (*spa*1,2,3; Laubinger et al., 2004), *phyA-219* (Reed et al., 1994), *phyB-*9 (Yoshida et al., 2018)*, cry1-304 (*Ahmad and Cashmore, 1993*), cry2-1* (Guo et al., 1998) and *phyBY276H* (YHB; Jones et al., 2015).

### Identification of the cry1M32 mutant

Based upon earlier research in our laboratory (Galvez-Valdivieso et al., 2009; Gorecka et al., 2014) in which we studied a possible role for heterotrimeric G protein-mediated high light (HL) signaling, we set out to identify candidate genes coding for 7 transmembrane proteins that may have a role as G protein coupled receptors. A collection of 59 T-DNA insertion mutants in genes coding for putative 7- transmembrane proteins (Moriyama et al., 2006; a kind gift from Professor Alan Jones, University of North Carolina) was screened for perturbed chlorophyll fluorescence in response to HL exposure (see below). The screening revealed that the insertion line Sail_1238_E12 (hereafter termed M32) was deficient in dynamic acclimation (Fig. 8B). The information available on T-DNA flanking sequences indicated that this was a T-DNA insertion in the first exon of At4g21570, a gene encoding a transmembrane protein of unknown function. However, complementation of M32 by transformation with the wild type At4g21570 gene did not restore a wild-type phenotype (data not shown).

Besides being defective in dynamic acclimation, M32 was impaired in blue light inhibition of hypocotyl elongation under both low and high blue light fluence, accumulated less chlorophylls and anthocyanins than Col-0 under blue light, and presented delayed flowering time when grown in short day photoperiod. (Supplemental Fig. 8 B-E). Later and in the light of our subsequent hypothesis that *CRY1*-mediated signaling controls dynamic acclimation in Arabidopsis (see Results and Discussion), we realized that M32 resembled the phenotype of known *cry1* mutants. Therefore, we tested if *CRY1* was altered in this mutant. *CRY1* was amplified from its genomic DNA and the PCR product was Sanger sequenced on both strands. Col-0 *CRY1* amplicon was also sequenced. The analysis of the sequence showed that in M32, *CRY1* contains a single point mutation (G→A), which caused a substitution of Gly_347_Arg mutation in CRY1 (Supplemental Fig. 8F). This mutation was previously identified in a screening of EMS-mutagenized Arabidopsis seedlings (Ahmad et al., 1995) and designated as *hy4-15*, and affects the domain comprising the photolyase signature sequence. As a consequence, *hy4-15* plants produce a wild type amount of full length CRY1, but the protein is not functional. Therefore, we concluded that M32 mutant is in fact a *cry1* mutant that we named *cry1M32*.

### HL exposures

The HL exposure was a PPFD of 1100 (± 100) µmol m^-2^ s^-1^ from a white light emitting diode (LED) array (Isolight 4000; Technologica Ltd, Colchester UK) as described previously (Gorecka et al., 2014) and which permitted the simultaneous exposure of 9 plants. The spectrum of the LED array is shown in Supplemental Figure 7. The HL exposure raised leaf temperature by 5 °C within 5 min of exposure which remained at this level for the remainder of the experiment (Gorecka et al., 2014). To determine the effect of this raised temperature (and the accompanying change in VPD) on the wider leaf transcriptome, we carried out a microarray analysis on plants exposed to HL for 30 mins, or 27 °C under LL for 30 mins (LL/27 °C) compared with LL/22 °C control plants. There were 609 DEGs (1.5-fold change; FDR < 0.05) that responded to HL and/or LL/27 °C (Supplemental Data Set 12; see also GSE87755 and GSE87756). Of these DEGs, 73 responded to the temperature increase alone (Supplemental Data Set 12). Given the small number, these were not eliminated from the time series data but none of the (co)TF DEGs (Supplemental Data Set 4) fell into this group.

For the HL time series transcriptomics, two consecutive sowings, 24h apart, were grown to 35 dpg on the same growth room shelf and randomized across the shelf every day. Leaf 7 (Boyes et al., 2001) was tagged at 30 dpg. We used this staging of plant growth and 3 LED Isolight arrays to treat 27 plants each day. The HL exposure began 1h after subjective dawn and was completed 1h before subjective dusk. Each set of tagged leaves (4) at each HL time point and their LL controls (4) were sampled within 5 min at time 0.5h and each 0.5h interval for the 6h exposure. Two HL experiments were conducted with duplicate samplings of a full range of time points on each day. In addition, 4 time zero samples were processed for the 0h time point. Both HL experiments provided a total of 100 samples for RNA extraction. These were 4 biological replicates (*i.e*. 4 sampled leaves) per timepoint per HL treatment (48 samples) and LL control (48 samples) plus 4 zero time point samples.

To elicit dynamic acclimation, plants were subjected to 4h HL, followed by a 0.5h dark adaptation and then exposed to a range of actinic PPFDs (over 50 min) to collect CF data (see below). This HL treatment was repeated daily and CF data collected from the same plants for 5 consecutive days or on Days 1 and 5 only as stated.

### CF measurements and imaging

During the time series HL experiments, CF measurements were taken from leaf 7 of one plant i*n situ* under each isolight using PAM-2000 portable modulated fluorimeters (PAM-2000, Walz GmbH, Effeltrich, Germany). At the end of each experiment the dark-adapted CF parameter Fv/Fm was determined for the same plants and LL controls and then again 24h after being returned to growth conditions.

For dynamic acclimation experiments, photosynthetic efficiency was estimated with a CF imaging system (Fluorimager, Technologica Ltd, Colchester, UK), exposing the plants to increasing actinic PPFD from 200–to-1400 µmol m^-2^ s^-1^ in 200 µmol m^-2^ s^-1^ steps every 5 min as described previously (Barbagallo et al., 2003; Gorecka et al., 2014). Whole rosette CF images were collected at each PPFD and processed using software (Technologica Ltd) to collect numerical data typically from fully expanded leaves (≥ 4 per plant) for Fq’/Fm’, Fv’/Fm’ and Fq’/Fv’ (Barbagallo et al., 2003; Baker, 2008, Gorecka et al., 2014). In some experiments, the diminished size of mutant plants rendered image processing problematic and in such stated cases, whole rosette data were collected. The raw data were fed *via* Excel into a program in R to calculate, plot and statistically analyse the CF parameters (Gorecka et al., 2014). The fluorimager software produces average data of all leaf pixel values. CF parameters were represented as mean ± SE from a minimum of 4 plants, and statistical significance was estimated with ANOVA followed by a *post-hoc* TukeyHSD test.

### Measurement of photosynthesis

*A* was measured on leaf 7 of plants at 49 dpg using an infrared gas exchange system (CIRAS-1, PP Systems, Amesbury, MA, USA). The response of *A* to changes in the intercellular CO_2_ concentration (*C_i_*) was measured under a saturating PPFD, provided by a combination of red and white LEDs (PP Systems, Amesbury, MA, USA). In addition, the response of *A* to changes in PPFD from saturating to sub-saturating levels was measured using the same light source at the current atmospheric CO_2_ concentration (390 µmol mol^-1^). All gas analysis was made at a leaf temperature of 20 (±1) °C and a VPD of 1 (±0.2) KPa. Plants were sampled between 1 and 4 hours after the beginning of the photoperiod. For each leaf, steady state rates of *A* at current atmospheric [CO_2_] were recorded at the beginning of each measurement.

### Relative ion leakage

The method described by Overmyer et al. (2008) was followed. Briefly, leaves were collected from plants and placed in 5 ml de-ionized water, incubated with rotary shaking (100 rpm) for 4h and the conductivity of the solution determined with a conductivity meter (Mettler Toledo, Leicester, UK) calibrated according to the manufacturer’s instructions. Leaves were frozen overnight, thawed and conductivity measured again. Relative ion leakage was expressed as ’conductivity after 4 h’ / ’conductivity after freeze-thawing’.

### RNA extraction, labelling and hybridisation to microarrays

For the time series HL experiment, RNA was extracted from leaf 7 samples, labelled and hybridised to CATMA (a <u>C</u>omplete <u>A</u>rabidopsis <u>T</u>ranscriptome <u>M</u>icro<u>A</u>rray) microarrays (v3; Sclep et al., 2007) as described by Breeze et al. (2011). Two technical replicates were used per biological replicate. Four biological replicates with a total of 13 time points per treatment (HL and LL) were analysed in this way, resulting in a highly replicated high-resolution time series of expression profiles. The hybridisation of labelled cDNA samples’ experimental design for the HL and LL time series followed a statistically randomised loop design (Supplemental Fig. 9), which enabled expression to be determined at different time points both within and between treatments. After hybridization and washing, microarrays were scanned for Cy3 and Cy5 fluorescence and analysed as below. The raw and processed data are deposited with NCBI GEO (GSE78251).

### Analysis of Microarray Data

This has been described in detail previously (Breeze et al., 2011; Windram et al., 2012). Briefly, a mixed model analysis using MAANOVA (Wu et al., 2003; Breeze et al., 2011) was used with the same random (dye, array slides) and fixed variables (time point, treatments and biological replicate) to test the interaction between these factors as for the analysis of time series microarray data for senescing, *Botrytis cinerea*-infected, *Pseudomonas syringae*-infected and drought-stressed leaves (Breeze et al., 2011; Windram et al., 2012; Lewis et al., 2015; Bechtold et al., 2016). Predicted means were calculated for each gene probe for each of the combinations of treatment, biological replicate and time point and for each of the combinations of treatment and time point from averages of the biological replicates.

A GP2S Bayes’ factor (Stegle et al., 2010) was used to rank probes and genes in order of likelihood of differential expression over the whole of the time series. Inspection of selected probes from the rank order of likelihood of differential expression was used to identify significant changes in expression with a Bayes’ factor cut-off >10 giving 4069 probes corresponding to 3844 DEGs (Supplemental Data Set 1).

### Clustering of Gene Expression Profiles

The expression patterns of the identified DEGs in HL and LL were co-clustered with SplineCluster (Heard et al., 2005), using the mean expression profiles of the biological replicates generated from MAANOVA and a prior precision value of 0.001 as described previously (Windram et al., 2012; Bechtold et al., 2016).

### GO analysis

GO annotation analysis was performed using DAVID (Huang et al., 2008) or AGRIGO (Du et al., 2010) with the GO Biological Process (BP) category (Ashburner et al., 2000). Overrepresented GO_BP categories were identified using a hypergeometric test with an FDR threshold of 0.05 compared against the whole annotated genome as the reference set.

### Comparisons with published transcriptomics data

The 3844 HL DEGs were compared on a cluster-by-cluster basis with publicly available transcriptomics data. The references for each dataset are in the main reference section of the paper. Each DEG list from published data was mapped to AGI codes when necessary, cleaned to obtain single AGI codes since in some microarray data, probes mapped to several genes or were listed as “no_match” and were eliminated from the list. Overlaps within each cluster and their statistical significance were determined using a Hypergeometric Distribution Test (phyper function in R (v3.2.1)) in a custom R script, available upon request. When required, Venn diagrams of overlaps between Data Sets were plotted with Venny (http://bioinfogp.cnb.csic.es/tools/venny/index.html) and the significance of the overlaps calculated using the R phyper function.

### VBSSM

A full description of VBSSM applied to this type of time series transcriptomics data is provided in Bechtold et al. (2016). The individual expression data for each biological replicate (n=4) for selected DEGs in HL was run through the VBSSM algorithm (Beal et al., 2005) on a local server at the University of Essex (Bechtold et al., 2016) to generate the GRNs as described in Results. The VBSSM output files were imported, mapped and plotted with Cytoscape (Shannon et al., 2003; http://www.cytoscape.org/).

### Expression profiling by RNAseq

Mature (non-senescent) leaves were excised from 4 biological replicate plants per treatment, their total RNA extracted, and quality controlled as previously described (Albihlal et al., 2018). Library construction after mRNA enrichment and double stranded cDNA synthesis carried out using Illumina protocols by Novogene (UK) Ltd (Cambridge, UK; en.novogene.com/). Library sequencing was carried out on an Illumina HiSeq 4000 with a 150 base paired end reads to a depth of 20 million. Extraction and quality control of data from raw fastq files was carried out using the program CASAVA (Hosseini et al.,2010). The mapping of reads to the TAIR10 Arabidopsis genome sequence, followed by sorting and indexing of BAM output files was carried out using default settings in the program HISAT2 (v2.0.5; Kim et al., 2015). Across all samples, > 92.5% of bases read attained the Q30 score threshold. Transcript assembly and quantification was as fragments per kilobase of transcript sequence per million base pairs sequenced (FPKM) using HTseq (in union mode; Anders et al., 2015). Determination of differential expression between different genotypes and treatments was done using the program DEseq2 (Love et al., 2014) after read count normalisation and an adjusted *p* value threshold of <0.05 (negative binomial distribution *p* value model and FDR correction). Raw and processed data files were deposited in NCBI Gene Expression Omnibus (GSE158898).

### Locus codes of genes mentioned in the paper

AT1G01720, *ATAF1*; AT1G04400, *CRY2*; AT1G06180, *MYB13*; AT1G09100, *AAA-ATPase*; AT1G09570, *PHYA*; AT1G14150, *PnsL2*; AT1G14920, *GAI*; AT1G16300, *GAPCP-2*; AT1G22190, *RAP2.4*; AT1G22640, *MYB3*; AT1G25540, *PFT1*; AT1G25550, *MYB-like*; AT1G29910, *Lhcb1/*CAB3; AT1G29920, *CAB2/LHCII*; AT1G29930, *CAB1/LHCII*; AT1G43670, *FBPASE*; AT1G44575, *PsbS*; AT1G49720, *ABF1*; AT1G50420, *SCL3*; AT1G50640, *ERF3*; AT1G68520, *BBX14*; AT1G69010, *BIM2*; AT1G69490, *NAP*; AT1G70000, *MYBD*; AT1G75540, *BBX21*; AT1G76100, *PETE1*; AT1G76570, *LHCB7*; AT1G77450, *NAC032*; AT1G79550, *PGK*; AT2G01290, *RPI2*; AT2G05070, *LHCB2*; AT2G18790, *PHYB*; AT2G21330, *FBA1*; AT2G24540, *AFR*; AT2G24790, *BBX4*; AT2G27510, *FD3*; AT2G28350, *ARF10*; AT2G30790, *PSBP-2*; AT2G32950, *COP1*; AT2G34430, *LHB1B1*; AT2G34720, *NF-YA4*; AT2G35940, *BLH1*; AT2G40100, *LHCB4.3*; AT2G40970, *MYBC1*; AT2G43010, *PIF4*; AT2G46270, *GBF3*; AT2G46340, *SPA1*; AT3G08940, *LHCB4.2*; AT3G09640, *APX2*; AT3G21150, *BBX32*; AT3G27690, *LHCB2.3*; AT3G60750, *TK*; AT3G61190, *BAP1*; AT4G05180, *PSBQ-2*; AT4G05390, *RFNR1*; AT4G08920, *CRY1*; AT4G10180, *DET1*; AT4G10340, *LHCB5*; AT4G15090, *FAR1*; AT4G17460, *HAT1*; AT4G29190, *OZF2*; AT4G32730, *PC-MYB1*; AT4G38960, *BBX19*; AT5G01600, *FER1*; AT5G07580, *ERF106*; AT5G08520, *MYBS2*; AT5G11260, *HY5*; AT5G11530, *EMF1*; AT5G12840, *NF-YA1*; AT5G15210, *HB30*; AT5G28450, *LHC1*; AT5G38420, *RBCS2B*; AT5G38430, *RBCS1B;* AT5G42520, *BPC6*; AT5G43270, *SPL2*; AT5G44190, *GLK2*; AT5G51190, *ERF105*; AT5G61270, *PIF7*; AT5G61590, *ERF107*; AT5G62000, *ARF2*; AT5G65310, *HB5*; AT5G67300, *MYBR1*; AT5G67420, *LBD37*; ATCG00020, *PSBA*; ATCG00270, *PSBD*; ATCG00300, *PSBZ*; ATCG00350, *PSAA*.

## SUPPLEMENTAL DATA SETS, TABLES AND FIGURES

**Supplemental Figure 1.** Examples of temporal differentially expressed clusters.

**Supplemental Figure 2.** A comparison of plants exposed to daily HL for 5 days with their equivalent age LL controls and induction of dynamic acclimation in older plants.

**Supplemental Figure 3.** The HL exposure used does not produce extensive photodamage to leaves.

**Supplemental Figure 4.** The first draft inferred HL gene regulatory network and dynamic acclimation in *lhy-21* plants.

**Supplemental Figure 5.** Temporal patterns of transcript abundance of the 12 most connected genes in the BBX32-centric GRN under LL and HL conditions

**Supplemental Figure 6.** Dynamic acclimation of *phyB-9*, *phyA-211*, *cry2-1* and *pif4-2* plants compared with Col-0.

**Supplemental Figure 7.** Spectral properties of the growth lights and isolight used for HL exposure.

**Supplemental Figure 8.** Identification and characterisation of *cry1*-M32.

**Supplemental Figure 9.** A randomized loop design employed for loading samples onto the two-channel CATMA arrays.

**Supplemental Data Set 1.** The 3844 HL DEGs on a cluster-by-cluster basis.

**Supplemental Data Set 2.** Significantly enriched Biological Process GO Terms for the HL DEGs in each temporal cluster.

**Supplemental Data Set 3.** Statistical data for CF measurements of 24 - 28 dpg Col0 plants and 44-48 dpg Col-0 plants undergoing dynamic acclimation.

**Supplemental Data Set 4.** Annotated transcription (co-)factor genes differentially expressed in HL vs. LL leaves.

**Supplemental Data Set 5.** Significant overlaps, on a cluster-by-cluster basis, between publicly available gene expression datasets and transcriptomic meta-analyses for mutants and treatments that perturb chloroplast-nucleus signalling, ROS-metabolism or are undergoing dynamic acclimation.

**Supplemental Data Set 6.** Photosynthetic efficiency of *BBX32*-OE, *bbx32-1*, *hy5-2* and *hy5-215* plants during induction of dynamic acclimation.

**Supplemental Data Set 7.** Transcripts that show a significant change in response to 3.5h HL in Col-0 and/or in *BBX32*-OE LL and HL plants.

**Supplemental Data Set 8.** FKPM values for transcripts coding for (A) photosynthesis-associated nuclear and plastid-encoded genes and (B) (co-)TFs in the inferred gene regulatory network in Figure 3.

**Supplemental Data Set 9.** Significantly enriched GO BP Terms common between *BBX32*-OE LL / Col-0 LL, *BBX32*-OE HL / Col-0 HL and HL/LL Col-0 DEGs.

**Supplemental Data Set 10.** DEGs in the GO BP terms common between BBX32-OE / Col-0 LL, *BBX32*-OE HL / Col-0 HL and from the HL time series clusters.

**Supplemental Data Set 11.** Significantly enriched GO BP terms for HL DEGs that overlap with seedling-expressed genes commonly regulated by *COP1*, *SPA* and *PIF* gene families.

**Supplemental Data Set 12.** Differentially expressed genes from leaf 7 of Arabidopsis subjected to a 30 min temperature rise from 22°C to 27°C under LL and HL conditions.

## AUTHOR CONTRIBUTIONS

PMM, UB, GGV and RAF carried out the HL time series experiments and RNA extractions. PMM, RAF and EJS conducted the CF measurements for HL responses and dynamic acclimation. MER carried out the RNA extractions and with EJS and PMM, conducted the RNAseq data analysis. TL designed the gas exchange measurements, which were performed by JSAM and PAD. CAP, LB, JDM, AM, RAF and PMM were responsible for microarray data generation and analysis. RAF and PMM conducted the VBSSM modelling under the guidance of CAP. PMM, KJD, DLW, JB, VB-W and UB developed the HL time series project. PMM conceived and wrote the manuscript with contributions from all authors.

## ACKNOWLEDGEMENTS

We thank Silvere Vialet-Chabrand and Steven Driever for discussions and advice on statistical and physiological aspects of the study. Laura Flanders and Rahjish Khanna for the *BBX32* OE lines and *bbx32-1*. We acknowledge European (Nottingham) Arabidopsis Stock Centre for providing *cry1-304*, *cry2-1*, *phyA-219*, *phyB-9*, *pifq*, *pif4-1*, *lhy-21*, and *det1-1*. Matthew Jones and Ute Hoecker respectively are gratefully acknowledged for providing seed of *cop1-4*, Col-0 YHB and *spa1/spa2/spa3*. This work was supported by the UK Biotechnology and Biological Sciences Research Council (grants BB/F005822/1 and BB/F005806/1). CAP and DLW were supported by the UK Engineering and Physical Science Research Council (grant EP/I036575/1). JSAM was supported by NERC-CASE award (ENV-EATR-DTP: NE/L002582/1).

---

**Supplemental Figure 1. Examples of temporal differentially expressed clusters.** These graphs are representative depictions of 6 of the 43 temporal clusters generated by SplineCluster (see Methods) from the 3844 DEGs identified from the HL and LL time series microarray data. The graphs are program outputs, but the lettering has been enhanced for clarity.

**Supplemental Figure 2. A comparison of plants exposed to daily HL for 5 days with their equivalent age LL controls and induction of dynamic acclimation in older plants.**

**(A)** The CF parameter Fq’/Fm’ and inset graphs of Fq’/Fv’ and Fv’/Fm’ of fully expanded leaves (≥ 4) from 4 Col-0 plants (24 - 28 dpg) through an incremental series of actinic PPFDs (from 200 - 1400 µmol m^-2^ s^-1^) as described in Methods and legend of Figure 2. The data were collected as CF images and processed digitally to collect values from expanded leaves. Prior to these measurements, the plants (5d HL; dashed lines) were exposed to HL for 4h per day for 5 days while control plants (5d LL; solid lines) were kept under LL conditions. CF data was collected on the fifth day after the final HL exposure. The data (means ± SE) are combined from 5 plants from a single experiment.

**(B)** The CF parameters Fq’/Fm’, Fq’/Fv’ and Fv’/Fm’ respectively of fully expanded leaves (≥ 4) from 4 plants (44 - 48 dpg) through an incremental series of actinic PPFDs (from 200 - 1400 µmol m^-2^ s^-1^) as described in Methods and legend of Figure 2. The data were collected and processed as in A. Prior to these measurements, the plants were exposed to HL for 4h per day for 5 days, CF data being collected each day for 5 days: day 1 (blue), day 2 (red), day 3 (olive green), day 4 (purple) and day 5 (light blue). The data (means ± SE) are combined from 6 plants from a single experiment. The differences between days 1 and 5 measurements from panels A and B were significant (P <0.05; ANOVA and TukeyHSD) for all CF parameters indicated by asterisks (*). A full statistical analysis is available in Supplemental Data Set 5.

**Supplemental Figure 3. The HL exposure used does not produce extensive photodamage to leaves.**

**(A)** Fv/Fm values in leaf 7 of the plants in (A) prior to HL exposure, just after 6h HL and 24h later, having been returned to growth conditions. Plants were dark adapted for 30 min and Fv/Fm determined by CF imaging (see Methods).

**(B)** Relative ion leakage (conductivity) in leaf 7 of the plants in (A) after the 6h HL measurements compared with the LL control.

**(C)** Expression of HL-responsive H_2_O_2_ marker genes (*APX2* and *FER1*; Karpinski et al., 1997; 1999; Dietz et al., 2002; Ball et al 2004; Davletova et al., 2004; Mittler et al., 2004; Rossel et al., 2007; Oelze et al., 2012; Jung et al., 2013; Shao et al., 2013) and ^1^O_2_ marker genes (*AAA-ATPase* and *BAP1*; Kim et al., 2012; Ramel et al., 2012; 2013; Shao et al., 2013). Log_2_-transformed fluorescence values (mean ± SE; n= 4) were normalised with respect to the same values at the zero time point, are shown for the HL (dotted line) and LL (solid line) datasets, and asterisks denote statistical differences (P < 0.05; Student’s t test) at each timepoint.

**Supplemental Figure 4. The first draft inferred gene regulatory network and dynamic acclimation in lhy-21 plants.**

**(A)** The network shown was generated from the time series expression data for HL DEGs. The networks were generated using VBSSM (threshold z-score = 2.33; see Methods) and visualised using Cytoscape (v3.3.0; Shannon et al., 2003). The network shown is the first iteration of the modelling, which included expression data for *LHY*. Those nodes coloured yellow highlight *LHY*, *BBX32* and *HY5*.

**(B)** PSII operating efficiency (Fq’/Fm’) of fully expanded leaves (≥ 4) from 4 Ws-0 and 4 *lhy21* plants (24 - 28 dpg) through an incremental series of actinic PPFDs (from 200 - 1400 µmol m^-2^ s^-1^) as described in Methods and legend of Figure 2. Prior to these measurements, the plants were exposed to HL for 4h per day for a total of 5 days and the CF images were collected at day 1 (black lines) and day 5 (red lines) of the daily HL treatments for *lhy21* (dashed lines) and Col-0 (solid lines). The determination of the parameters was by CF imaging as described in the legend of Figure 2 and Methods. The data (means ± SE) are combined from 5 plants from a are nodes of ≥ 3 edges (connections) in the GRN shown in Figure 3. The data are from the microarray time series transcriptomics (Fig.1A; Supplemental Data Set 1). The numbers in parentheses is the temporal cluster to which each gene was assigned (Fig. 1; Supplemental Data Set 1). The asterisk denotes a significant difference at that time point between LL and HL samples (P ≤ 0.05; ANOVA).

**Supplemental Figure 6. Dynamic acclimation of *phyB-9*, *phyA-211*, *cry2-1* and *pif4-2* plants compared with Col-0.**

The plots show the PSII operating efficiencies (Fq’/Fm’) determined from CF images of ≥ 4 mature leaves from 8 plants (24-28 dpg) from two experiments (means ± SE) in (A-C) and a single experiment of 4 plants in (D). The plants had been exposed to 4h HL each day for 5 consecutive days (see Methods and legend of Figure 2). CF parameter values were collected at a range of actinic PPFDs (as indicated on the x-axis) at the end of days 1 and 5 of HL. The Fq’/Fm’ values at day 1 (black lines) and day 5 (red lines) for mutant plants (dashed line) and Col-0 (solid line) of the HL treatments for (A) *phyB-9*, (B) *phyA-211*, (C) *cry2-1* and (D) *pif4-2.* All CF data points comparing Col-0 and the mutants were not significantly different (P > 0.1; ANOVA and TukeyHSD).

**Supplemental Figure 7. Emission spectra for the growth lights and for the LED-isolight used to induced dynamic acclimation.**

**Supplemental Figure 8. Identification and characterisation of *cry1*-M32.**

**(A)** Dark-adapted Fv/Fm measurements on Arabidopsis 2 weeks-old seedling taken by chlorophyll fluorescence imaging (see Methods). Plants (n=15) were measured before HL exposure (LL), after 5-fold HL exposure for 1 hour. Values are the means ± SE

**(B)** Fluence response curves of 6-d-old Col-0 and M32 mutant seedlings grown in blue light.

**(C)** (Effect of day length on flowering time. Chlorophylls (C) and anthocyanins.

**(D)** Chlorophyll content of 8 d.p.g. Col-0 and M32 mutant seedlings grown in continuous BL (40 µmol·m^-2^·s^-1^). Error bars represent SE.

**(E)** Anthocyanin content of the same seedlings as in (D).

**(F)** Mutation of M32 at the photolyase signature sequence of CRY1.

**Supplemental Figure 9. A randomized loop design employed for loading samples onto the two-channel CATMA arrays.**

A randomised loop design was employed for loading samples onto the two-channel CATMA arrays, enabling expression analysis between timepoints (measured in hours of treatment under LL or HL), both within and between treatments. (A) shows the within-treatment samples that were loaded on the same array, but differentially labelled with either Cy3 or Cy5, and (B) shows the cDNA samples from the two different treatments that were differentially labelled and loaded onto the same array.

